# Functional and dysfunctional T regulatory cell states in human tissues in RA and other autoimmune arthritic diseases

**DOI:** 10.1101/2025.09.23.677874

**Authors:** Byunghee Koh, Shani T. Gal Oz, Ryota Sato, Hung N. Nguyen, Garrett Dunlap, Chrisopher Mahony, Chrissy Bolton, Accelerating Medicines Partnership (AMP) RA/SLE Network, Lucy R. Wedderburn, Adam P. Croft, Laura Donlin, Soumya Raychaudhuri, Ilya Korsunsky, Deepak A. Rao, Michael B. Brenner

## Abstract

Regulatory T cells (Tregs), characterized by FOXP3 expression, are essential for maintaining immune homeostasis by controlling inflammation. However, in autoimmune diseases such as rheumatoid arthritis (RA), impaired Treg function contributes to immune dysregulation and disease pathology. While most studies of human Tregs have focused on blood, here we analyzed Tregs in synovial tissues from RA patients using single cell RNA sequencing (scRNAseq). We identified two predominant Treg states, CD25^hi^CXCR6^pos^ Tregs with strong suppressive function, and CD25^lo^AREG^pos^ Tregs, a dysfunctional state exclusively enriched in synovial tissues but not in blood. Computational and in vitro analyses revealed that cortisol induced AREG expression, suppressed glycolysis, and impaired the suppressive function of CD25^lo^AREG^pos^ Tregs. In turn, AREG promoted an IL-33^+^ inflammatory phenotype in synovial fibroblasts. Importantly, we found that TNFR2 engagement can prevent or reverse this dysfunctional Treg state. In contrast to CD25^lo^AREG^pos^ Tregs, CD25^hi^CXCR6^pos^ Tregs were highly suppressive, showed coordinated abundance with macrophages in synovial tissue, and functionally interacted with membrane-bound TNFα expressed by macrophages, which promoted their functional suppressive state. These two Treg subsets were similarly found in the synovial tissue in Juvenile Idiopathic Arthritis (JIA), another inflammatory arthritic disorder, indicating conserved mechanisms across arthritic diseases. Together, our findings define distinct pathways driving divergent functional and dysfunctional Treg states in inflamed tissues and point to interventions that may prevent or reverse the development of the dysfunctional state.

## Introduction

Rheumatoid arthritis (RA) is a chronic autoimmune disease characterized by persistent inflammation in synovial tissues leading to degradation of joint cartilage and bone ^1,2^. This inflammatory process is driven by infiltrating immune cells including lymphocytes and monocytes, which produce pro-inflammatory cytokines such as tumor necrosis factor (TNF), interleukin-1β (IL-1β), and interleukin-6 (IL-6) ^3^. Regulatory T cells (Tregs), a specialized subset of CD4^+^ T cells, are essential for maintaining immune tolerance and suppressing excessive immune responses ^4,5^. Tregs are defined by high expression of CD25, and the transcription factor FOXP3, both of which are crucial for their development, stability, and function ^6^. CD25 enables Tregs to respond to low levels of IL-2, a cytokine critical for their development and function ^7^.

Recent studies have highlighted the heterogeneity of Tregs across tissues and disease states. In non-lymphoid tissues, Tregs can differentiate into specialized subsets that adapt to local environmental cues, including cytokines and chemokines, and acquire distinct phenotypes suited for their functions at specific inflammatory sites ^8–12^. This functional and phenotypic diversity reflects the varied roles that Tregs play in regulating immune responses and interacting with different stromal and immune cell populations.

In RA, Tregs migrate to the synovium to counteract inflammation and limit tissue damage caused by pathogenic immune cells ^13–15^. However, the inflamed synovial microenvironment may compromise Treg stability and suppressive function. For instance, IL-6 produced by synovial fibroblasts can drive the conversion of Tregs into pro-inflammatory Th17 cells ^16^, while IL-1β promotes osteoclastogenic Treg differentiation ^17^. Soluble TNFα (sTNFα) impairs Treg function through TNFR1-mediated FOXP3 dephosphorylation ^18^, whereas membrane-bound TNFα (mTNFα) signals through TNFR2 to stabilize FOXP3 and maintain Treg suppressive capacity ^19,20^. Although sTNFα can bind both receptors, mTNFα more efficiently activates TNFR2 ^21^. In murine models, Treg-specific TNFR2 deficiency leads to impaired Treg accumulation and function, exacerbating disease in multiple autoimmunity models ^19,22^. In human Tregs, TNFR2 stimulation enhances proliferation, stability, and function via the non-canonical NF-κB pathway and increased glycolysis via the PI3K-mTOR axis ^23,24^.

Metabolic adaptation is increasingly recognized as a key determinant of Treg function. In lipid-rich environments such as visceral adipose tissue, Tregs rely on fatty acid oxidation for oxidative phosphorylation (OXPHOS) ^25^, while in tumors, Tregs adapt to a glucose-depleted environment by using lactate for OXPHOS ^26^. Human Tregs exhibit greater reliance on glycolysis ex vivo than their murine counterparts, which is critical for both their suppressive function^27^ and migratory capacity ^28^. However, the metabolic state of Tregs in chronically inflamed human tissues such as RA synovium remains poorly understood.

Most studies characterizing human Tregs in RA have focused on cells from peripheral blood and synovial fluid of RA patients. However, the synovial tissue itself represents a distinct chronic inflammatory niche, with a unique cellular composition including activated synovial fibroblasts and tissue-resident macrophages ^29,30^. This microenvironment produces a complex cytokine and chemokine milieu ^31^, which likely contributes to the shaping of tissue-adapted Treg phenotypes.

Here, we employed unbiased single cell RNA-seq (scRNAseq) analysis of Tregs isolated from RA synovial tissue and compared them to Tregs from synovial fluid and peripheral blood. We identified two major Treg subsets in diseased tissue; 1) CD25^hi^CXCR6^pos^ Tregs, which exhibit potent suppressive activity, and 2) CD25^lo^AREG^pos^ Tregs, which display reduced suppressive function and are uniquely enriched in synovial tissue. We demonstrated that the inflammatory synovial environment, particularly through fibroblast-derived glucocorticoids, drives the emergence of dysfunctional CD25^lo^AREG^pos^ Tregs mechanistically via metabolic reprogramming and AREG induction. In contrast, macrophage-derived mTNFα supports the expansion and function of functionally suppressive CD25^hi^CXCR6^pos^ Tregs. Finally, we identified similar Treg phenotypes in other arthritic diseases, suggesting conserved mechanisms across inflammatory arthritic conditions. Together, our findings provide a detailed map of functional and dysfunctional Treg states in inflamed human synovial tissue, reveal tissue-specific cues that govern their fate, and highlight therapeutic strategies to restore Treg function in chronic inflammation.

## Results

### Transcriptional heterogeneity in T regulatory cell states in RA tissues

In order to delineate the nature of Tregs in inflamed tissues from patients with RA, we conducted a single-cell analysis across synovial tissue, synovial fluid, and blood, utilizing four scRNAseq datasets (Datasets A-D, **Supplementary Table 1, Fig 1a**). We analyzed each dataset independently to identify Treg clusters based canonical Treg markers (*FOXP3*, *IL2RA* (encoding CD25)) in CD4^+^ T cells clusters (**Extended Data Fig. 1a-b, Supplementary Table 1**). The proportion of Tregs in RA synovial tissue, as calculated based on single cell proportions in dataset A, ranged between 3-24% of CD4^+^ T cells (average 10.2%, SD 4.6%; **Extended Data Fig. 1c, left**), which is consistent with flow cytometric analysis (**Extended Data Fig. 1c. right**). After selecting *FOXP3*-expressing cells in each dataset, we integrated 18,909 Tregs from synovial tissue, synovial fluid, blood from RA patients and blood from healthy donors (HD) and visualized their distribution using Uniform Manifold Approximation and Projection (UMAP) (**Fig. 1b**). Tregs showed a spectrum of *IL2RA* expression, broadly separating into CD25^high^ and CD25^low^ groups, as previously reported ^32^ (**Fig. 1c**). The expression of other canonical Treg markers, *FOXP3* and *IKZF2* (encoding Helios), varied in parallel with *IL2RA* expression (**Fig. 1d**).

**Figure 1.**
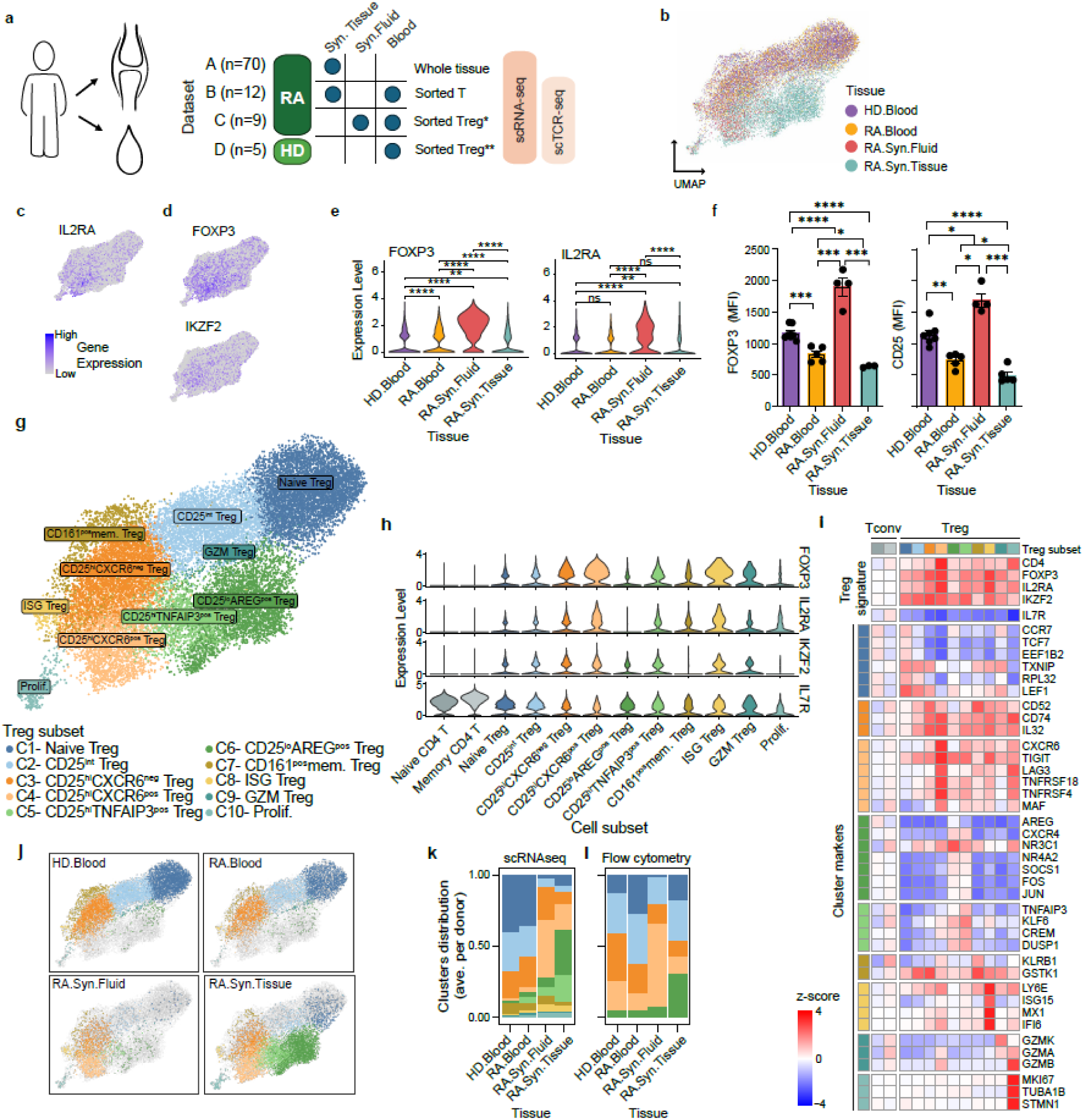
Single cell analysis of regulatory T cell subsets across tissues in rheumatoid arthritis. **a,** Schematic overview of the scRNA-seq study design. Samples were obtained from synovial tissue (datasets A-B ^32,75^), synovial fluid (dataset C, current paper) and blood (datasets B-C) from RA patients, and blood from healthy donors (HD) (dataset D, ^76^), and their corresponding TCR repertoire sequencing data (datasets B-D). Tregs were independently identified in each dataset and integrated using harmony ^79^. Clustering and downstream analyses were performed on 18,909 Tregs from all tissues. *Sorted Tregs: CD3^+^CD4^+^CD127^−^; **Sorted Tregs: CD4^+^CD25^+^CD127^−^. **b**, Uniform Manifold Approximation and Projection (UMAP) visualization of integrated Tregs from synovial tissue, synovial fluid, and blood (RA and HD), colored by tissue of origin. **c-d,** Expression of canonical Treg markers shown on UMAP: **(c)** *IL2RA* (CD25), **(d)** *FOXP3*, and *IKZF2* (HELIOS). Color scale ranges from grey (no expression) to dark purple (highest expression). **e,** Violin plots of *FOXP3* (top) and *IL2RA* (bottom) expression across tissues. Comparisons were performed using the Wilcoxon test. **f**, Flow cytometric analysis of FOXP3 and CD25 expression in Tregs across tissues (n=5). **g,** UMAP plot showing clustering of integrated Tregs, colored by cluster identity. **h,** Violin plots of canonical Treg markers: *FOXP3*, *IL2RA*, *IKZF2*; and the non-Treg marker *IL7R* (CD127) across 10 integrated Treg clusters and naïve/ memory CD4^+^ T cells from RA synovial tissue. **i,** Heatmap of pseudobulk expression for selected marker genes across Treg clusters and naïve/memory CD4^+^ T cells from RA synovial tissue. Marker genes were chosen based on cluster differential expression analysis and known Treg markers (separated by row blocks). Single-cell counts were summed into pseudobulk samples and log normalized for each cluster. Rows are z-scored-normalized relative to non-Treg clusters. Color represents the relative gene expression (blue: low; red: high). **j**, UMAP plots showing the distribution of Treg clusters across individual tissues. Cells from the focal tissue are colored by cluster; other cells are shown in grey. **k,** Proportions of Treg clusters in each tissue, calculated per donor and averaged. **l,** Flow cytometric analysis of Treg cluster proportion across tissues. Data represent mean ± SEM. Statistical significance was assessed using unpaired t-tests: \**p* < 0.05, \*\**p*<0.01, \*\*\**p* <0.001, \*\*\*\**p* <0.0001.

When comparing Tregs across RA tissues, we noticed higher *FOXP3* and *IL2RA* expression in synovial fluid compared to synovial tissue or blood (RA or HD) (**Fig. 1e**). We confirmed this difference at the protein level using flow cytometry (**Fig. 1f**). Louvain clustering identified 10 distinct Treg clusters (C1-C10) **(Fig. 1g**). All Treg clusters expressed elevated *FOXP3*, *IL2RA*, and *IKZF2* and reduced *IL7R* (CD127) compared to conventional naïve and memory CD4^+^ T cells, though expression levels varied across Treg clusters (**Fig. 1h**).

We annotated the clusters (C1-10) based on differential gene expression (**Fig. 1i**, **Extended Data Fig. 1d, Supplementary Table 1**). We focused subsequent analyses on the larger clusters (C1-C6). Smaller clusters (C7-C10) were designated as CD161^pos^ memory Tregs, ISG Tregs with enhanced interferon-stimulated genes expression, granzyme-expressing Tregs (GZM Tregs), and proliferating Tregs (Prolif.), respectively (**Fig. 1i**). C1 (naïve Tregs) exhibited enrichment in naïve T cell markers *CCR7* and *TCF7* with low *IL2RA*. C2 (CD25^int^ Treg) had intermediate *IL2RA* expression. C3 (CD25^hi^CXCR6^neg^ Treg) and C4 (CD25^hi^CXCR6^pos^ Treg) exhibited high *IL2RA* levels (**Fig. 1h-i**). C4 expressed elevated levels of genes associated with suppression, such as *TNFRSF4* (OX40) and *TNFRSF18* (GITR), along with *CXCR6*, a chemokine receptor associated with tissue residency and homing to inflamed sites ^33–35^. C5 and C6 expressed lower *IL2RA* and *FOXP3* expression but upregulated memory-associated genes including *JUN*, *JUNB*, *FOS*, and *KLF6* ^36^. Both clusters also expressed suppressor of cytokine signaling 1 (*SOCS1*) and *SOCS3*, negative regulators of STAT1 and STAT3, respectively ^37,38^. Importantly, C5 (CD25^lo^AREG^pos^ Treg) uniquely expressed high level of *AREG*, an epidermal growth factor receptor (EGFR) ligand implicated in tissue repair ^39–42^. C6 (CD25^hi^TNFAIP3^pos^ Treg) expressed high level of TNFα induced protein 3 (TNFAIP3, also known as A20), a critical negative regulator of inflammation linked to RA susceptibility ^43^.

Treg cluster proportions varied significantly across tissues (**Fig. 1j**). Blood from RA patients and healthy donors was dominated by naïve Tregs (C1), CD25^int^ Tregs (C2) and CD25^hi^CXCR6^neg^ Tregs (C3), while CD25^hi^CXCR6^pos^ Tregs (C4) were significantly enriched in synovial tissue and fluid compared to blood (FDR < 0.001, mean log2 fold enrichment = 3.6). In contrast, naïve Tregs (C1) and CD25^int^ Tregs (C2) were more abundant in blood (FDR<0.001 mean log2 fold enrichment = 2.8). Notably, CD25^lo^AREG^pos^ Tregs (C6) and CD25^hi^TNFAIP3^pos^ Tregs (C5) were predominantly found in synovial tissue (**Fig. 1j-k, Extended Data Fig. 1e)**. Flow cytometric validation confirmed these patterns, showing CD25^hi^CXCR6^pos^ Tregs enriched in synovial fluid and tissues, and CD25^lo^AREG^pos^ Tregs almost exclusively present in synovial tissue (**Fig. 1l, Extended Data Fig. 1f**).

Together, these findings revealed distinct transcriptional Treg states with tissue-specific distribution. CD25^lo^AREG^pos^ Tregs appear to be uniquely adapted or reprogrammed by the inflamed synovial microenvironment, whereas CD25^hi^CXCR6^pos^ Tregs display an activated, suppressive phenotype that distinguishes them from CD25^hi^CXCR6^neg^ Tregs and are enriched in synovial tissue and fluid. The reciprocal emergence of CD25^hi^CXCR6^pos^ Treg and CD25^lo^AREG^pos^ Treg states may reflect divergent differentiation paths of Tregs in inflamed tissue and provide insight into how local cues shape functional versus dysfunctional Treg responses in RA.

### Distinct clonal and functional states define CD25^hi^CXCR6^pos^ Tregs and CD25^lo^AREG^pos^ Tregs

To gain insight into the clonal architecture and developmental relationships among Treg subsets, we analyzed paired T cell receptor (TCR) and transcriptomic data from datasets B–D, spanning synovial tissue, synovial fluid, and blood. TCR “clones” were defined by identical nucleotide sequences of the Complementarity-Determining Region 3 (CDR3) in both the α and β chains within individual donors. Shared TCR signatures among clusters can indicate common developmental origins. We observed variable degrees of clonal expansion across Treg clusters (**Fig. 2a**), which positively correlated with *IL2RA* expression (**Fig. 1c**). CD25^hi^CXCR6^pos^ Tregs were the most clonally expanded subset overall, with 32% of cells sharing a clone (>1 cell) (**Fig. 2b**). Among tissues, Tregs from synovial fluid exhibited the highest degree of clonal expansion (**Fig. 2c, Extended Data Fig. 2a**). CD25^hi^CXCR6^pos^ Tregs were also the most clonally expanded subset in both synovial tissue (20%) and synovial fluid (44%). By comparison, ISG Tregs in RA and healthy donor blood showed similar expansion levels (**Extended Data Fig. 2b**). CD25^hi^CXCR6^neg^ Tregs and CD25^hi^TNFAIP3^pos^ Tregs were less clonally expanded, while CD25^int^ Tregs and CD25^lo^AREG^pos^ Tregs had the lowest expansion among non-naïve subsets. These results support the concept of increased proliferation among activated effector Tregs, particularly in synovial fluid.

**Figure 2.**
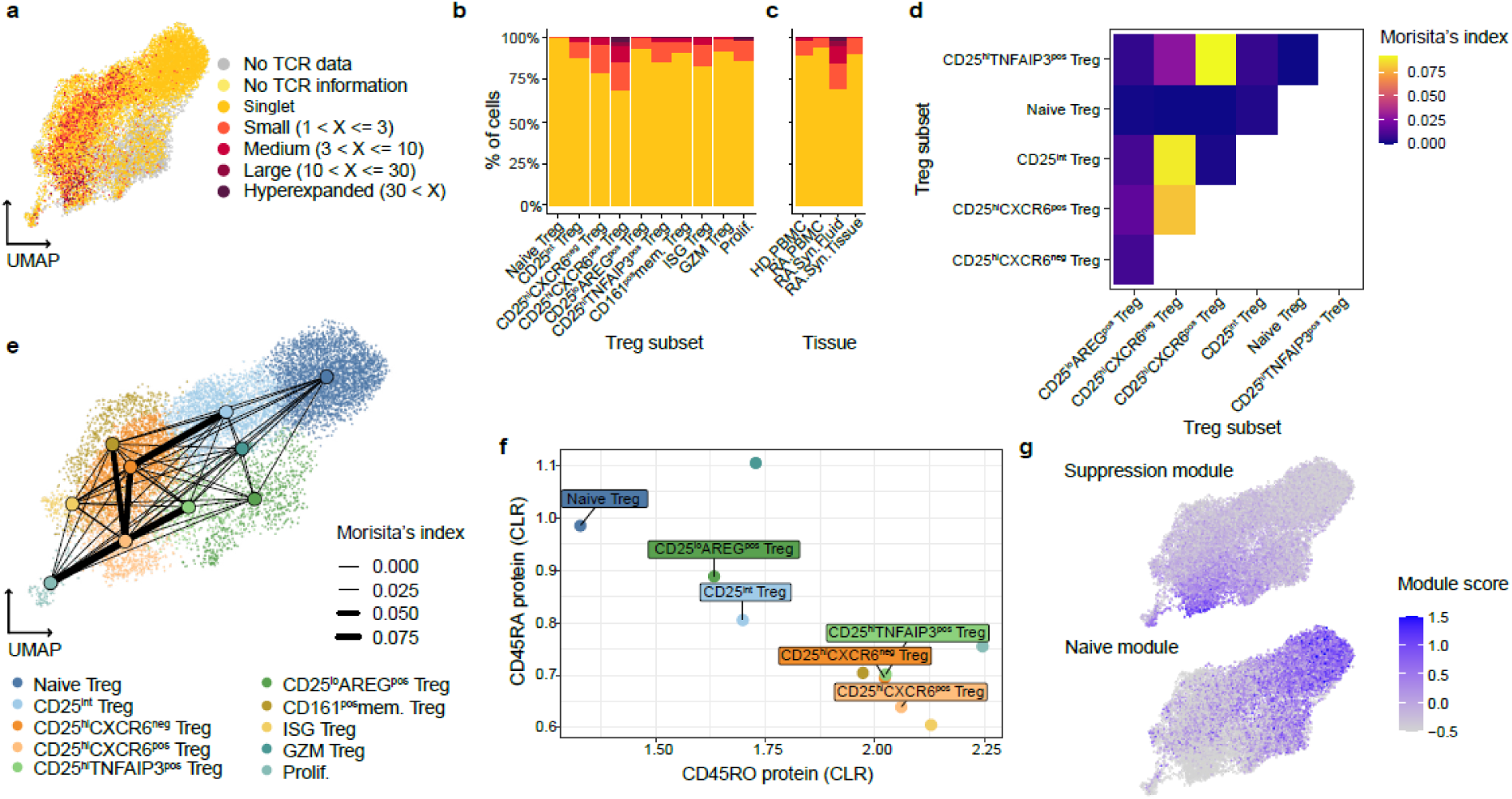
Clonal expansion and functional characteristics of Treg subsets. **a,** Clonal expansion visualized on a UMAP plot based on single cell TCR repertoire sequencing (scTCRseq) data. Cells are color-coded based on their clonal expansion level, determined by th CDR3 sequences of both α and β chains. “No TCR data” represents cells from dataset A, which lacked scTCRseq data. “No TCR information” represents cells with scRNAseq but were not captured by scTCRseq. Singlets denote cells with unique clones. Expanded clones are categorized as follows: Small (2-3 cells per clone), Medium (4-10 cells per clone), Large (11-30 cells per clone), and Hyperexpanded (>30 cells per clone). **b-c,** Proportions of clonally expanded cells in each Treg cluster **(b)** and in each tissue **(c)**. In both panels, only datasets B-D (with TCR data) are shown. Clonal categories match panel a. **d,** Heatmap of pairwise clonal overlap between selected Treg subsets, quantified using Morisita’s index. Higher values indicate greater clonal similarity. **e,** Clonal overlaps projected onto UMAP space. Cluster centroids are connected by lines, with line thickness representing the degree of clonal overlap (Morisita’s index). **f,** Average CLR-normalized protein expression levels (CITE-seq) of CD45RA (y-axis) and CD45RO (x-axis) for each Treg subset. Selected subsets are labeled. **g,** UMAP plots showing gene module scores for: Suppression module (left), naïve module (right). Color intensity represents the relative expression score for each module, with darker colors indicating higher expression.

To examine potential differentiation relationships among Treg subsets, we assessed shared TCR clonotypes between clusters using Morisita’s index (MI) as a measure of clonal overlap (**Extended Data Fig. 2c**). Among the larger clusters, we found that CD25^hi^CXCR6^pos^ Tregs showed the highest clonal overlap with CD25^hi^TNFAIP3^pos^ Tregs (MI = 0.091; 13 shared clones) and CD25^hi^CXCR6^neg^ Tregs (MI = 0.079; 33 shared clones) (**Fig. 2d, Extended Data Fig. 2d**). Similarly, CD25^int^ Tregs exhibited substantial overlap with CD25^hi^CXCR6^neg^ Tregs (MI=0.088, 102 shared clones). These relationships were visualized on a UMAP embedding, highlighting potential differentiation trajectories (**Fig. 2e**). Consistent with these MI scores, we observed the highest number of shared clonotypes between CD25^hi^CXCR6^pos^ Tregs and both CD25^hi^CXCR6^neg^ Tregs and CD25^hi^TNFAIP3^pos^ Tregs suggesting that CD25^hi^CXCR6^pos^ Tregs may arise from either or both two subsets (**Extended Data Fig. 2d**).

To further investigate the differentiation states of Treg subsets, we analyzed CD45RA/CD45RO protein expression using Cellular Indexing of Transcriptomes and Epitopes (CITE-seq) data from synovial tissue (Dataset A) (**Fig. 2f**). Naïve Tregs exhibited high CD45RA and low CD45RO expression, consistent with their transcriptional profile. In contrast, CD25^hi^CXCR6^pos^ Tregs and CD25^hi^CXCR6^neg^ Tregs exhibited the opposite patterns, consistent with their memory phenotype. CD25^int^ Tregs and CD25^lo^AREG^pos^ Tregs displayed intermediate CD45RA/CD45RO levels, suggesting transitional or early memory states. We next evaluated two gene expression modules: 1) a suppressive module (*ENTPD1, ICOS, TNFRSF9, IL10, TNFRSF4, TGFB1, TIGIT, CTLA4, TNFRSF18, LAG3*), and 2) a naïve module (*TCF7, CCR7, LEF1, SELL*) (**Fig. 2g**). Clonal expansion was positively correlated with the suppressive module score and negatively correlated with the naïve module score (**Fig. 2a, g**). Notably, CD25^lo^AREG^pos^ Tregs exhibited the lowest suppressive score, and highest naïve score among non-naïve subsets, suggesting CD25^lo^AREG^pos^ Tregs may derive from naïve Tregs but failed to fully acquire suppressive and memory phenotypes.

Together, these data highlight a key functional distinction between CD25^hi^CXCR6^pos^ Tregs, which are clonally expanded, suppressive, and memory-like, and CD25^lo^AREG^pos^ Tregs, which are less expanded, less suppressive, and exhibit features of a naïve or transitional state. These differences provide insight into the developmental and functional plasticity of Tregs within the inflamed RA synovial environment.

### CD25^hi^CXCR6^pos^ Tregs are highly suppressive Tregs in RA synovium

To elucidate functional differences among Treg subsets, we investigated their capacity to suppress conventional T cell proliferation. Gene expression analysis revealed that CD25^hi^CXCR6^pos^ Tregs expressed the highest levels of multiple suppressive molecules, including *CTLA4*, *ICOS*, *TNFRSF9*, *TNFRSF4*, *TNFRSF18*, *ENTPD1*, *TIGIT*, and *LAG3* (**Fig. 3a, Extended Data Fig. 3a**). Flow cytometry validation confirmed elevated expression of CTLA4 and ICOS in CD25^hi^CXCR6^pos^ Tregs (CXCR6^pos^) compared to other Treg subsets (CXCR6^neg^) (**Fig. 3b**). Functional validation using in vitro suppression assays showed that while both CD25^hi^CXCR6^pos^ and CD25^hi^CXCR6^neg^ Tregs from synovial fluid suppressed CD4^+^ conventional T cells (Tconv) proliferation, the CD25^hi^CXCR6^pos^ Tregs demonstrated superior suppressive capacity (**Fig. 3c, Extended Data Fig. 3b**).

**Figure 3.**
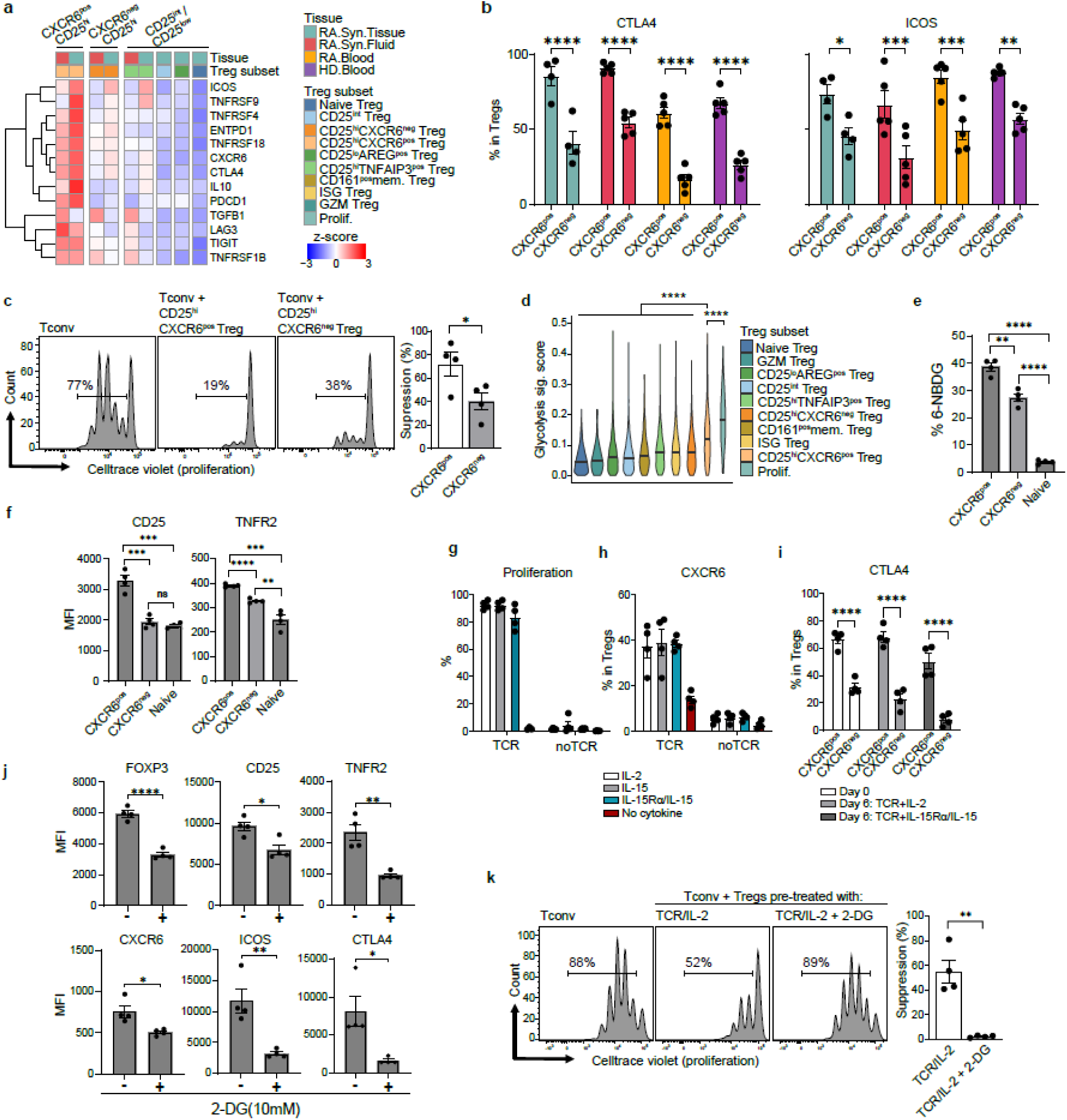
CD25^hi^CXCR6^pos^ Tregs exhibit enhanced suppressive capacity in RA synovium. **a,** Heatmap depicting Treg suppressive marker gene expression across subsets and tissues. Row-wise Z-scores of the normalized expression of Treg suppressive genes are shown. Single cell counts were summed into pseudobulk samples and log normalized for each tissue (top color bar) and Treg subset (bottom color bar). Only RA synovial tissue and synovial fluid cells are displayed. Treg subsets with less than 5% of cells from synovial fluid are shown only for synovial tissue Tregs. **b**, Flow cytometric analysis of suppressive markers in Tregs across tissues (n=5 donors). **c,** Suppressive capacity of CD25^hi^CXCR6^pos^ Tregs and CD25^hi^CXCR6^neg^ Tregs, isolated along with CD4^+^ Tconv from RA synovial fluid (n=4 donors). Left: representative histogram; right: mean suppressive capacity. Suppression (%) calculated as: 100 X [1-(percentage of dividing Tconv with Tregs/ percentage of dividing Tconv without Tregs). **d,** Glycolysis module score (based on 80 genes from the “HALLMARK_GLYCOLYSIS” gene set, MsigDB ^44,87^) shown across Treg subsets. The Wilcoxon test was used to assess differences in glycolysis module score between CD25^hi^CXCR6^pos^ Tregs and each of the other Treg subsets. **e-f,** Flow cytometric analysis of 6-NBDG uptake (e), and CD25 and TNFR2 expression (f) in indicated Tregs from healthy donor blood (n=4 donors). **g-i**, Effects of IL-2, IL-15 or IL-15/IL-15Rα complex on proliferation (g), CXCR6 (h) and CTLA4 (i) expression in Tregs stimulated for 5 days (n=4 donors). **j-k,** Effects of glycolysis inhibition by 2-DG on Treg phenotypes (j) and suppressive function (k) using same suppression assay as in (c). Data represent mean ± SEM. Statistical significances were determined by unpaired t-tests: \**p* < 0.05, \*\**p*<0.01, \*\*\**p* <0.001, \*\*\*\**p* <0.0001.

We next explored pathways that may promote the differentiation of CD25^hi^CXCR6^pos^ Tregs, focusing on metabolic pathways regulation. Using gene sets from Molecular Signatures Database (MsigDB) ^44^, we calculated module scores to assign a module score for each pathway. We assigned module scores for glycolysis (**Fig. 3d**), and oxidative phosphorylation (OXPHOS) (**Extended Data Fig. 3c**). CD25^hi^CXCR6^pos^ Tregs and proliferating Tregs exhibited the highest glycolytic and OXPHOS gene expression, followed by CD25^hi^CXCR6^neg^ Tregs and CD25^int^ Tregs, while naïve Tregs showed the lowest metabolic activity. To validate these findings, we isolated CD25^hi^CXCR6^pos^ Tregs, CXCR6^neg^ Tregs (CD25^hi^CXCR6^neg^ Tregs and CD25^int^ Tregs), and naïve Tregs from healthy donor blood and assessed glucose uptake via 6-NBDG (**Fig. 3e**). Consistent with their increased expression of glycolytic genes, CD25^hi^CXCR6^pos^ Tregs demonstrated the highest glucose uptake rate, followed by CXCR6^neg^ Tregs and naïve Tregs.

Given the link between metabolism and Treg function, we examined signaling pathways known to regulate glycolysis in Tregs, specifically IL-2/STAT5, IL-15 and TNFR2 signaling ^24,45^. CD25^hi^CXCR6^pos^ Tregs expressed higher levels of CD25 and TNFR2 compared to other Treg subsets in both healthy donor blood (**Fig. 3f**) and RA synovial tissues (**Extended Data Fig. 3d**). We then investigated the role of IL-2 and IL-15-both of which signal through IL-2Rβ and γ chain-in supporting CD25^hi^CXCR6^pos^ Treg differentiation. Because Tregs lack IL-15Rα, IL-15 requires trans-presentation via IL-15Rα on accessory cells for high-affinity signaling ^46^. Using plate-bound IL-15Rα conjugated with IL-15 (IL-15Rα/IL-15 complexes), we found that both TCR engagement and IL-2 or IL-15 were required for Treg proliferation (**Fig. 3g**) and CXCR6 expression (**Fig. 3h**). *In vitro* expanded CD25^hi^CXCR6^pos^ Tregs also expressed higher CTLA4 levels than CD25^hi^CXCR6^neg^ Tregs (**Fig. 3i**), consistent with their stronger suppressive phenotype. Phospho-STAT5 assay confirmed that Tregs can respond to high dose of IL-15, though less robustly than to IL-2 (**Extended Data Fig. 3e**). These findings suggest that in IL-15 rich environments, Treg can directly respond to IL-15, but in IL-15-limited environment such as RA synovium, IL-15Rα trans-presentation may be essential for optimal Treg function and homeostasis.

To test whether glycolysis is necessary for CD25^hi^CXCR6^pos^ Treg differentiation, we treated Tregs with 2-deoxy-D-glucose (2-DG), a glucose analog that inhibits glycolysis. 2-DG treatment reduced the expression of key markers of CD25^hi^CXCR6^pos^ Tregs, including FOXP3, CD25, CTLA4 and ICOS - consistent with previous studies ^24,27^-as well as CXCR6 and TNFR2 (**Fig. 3j**). Correspondingly, glycolysis inhibition impaired Treg suppressive function (**Fig. 3k**), indicating that glycolytic metabolism is essential for maintaining the phenotype and function of CD25^hi^CXCR6^pos^ Tregs.

Together, these findings show that enhanced glycolytic activity supports the differentiation and maintenance of the CD25^hi^CXCR6^pos^ Tregs in RA synovium. This phenotype is likely sustained through IL-2/STAT5, IL-15/STAT5, and TNFR2 signaling pathways. The specific contribution of TNFR2 to this process is addressed in the next section.

### Glucocorticoid and TNFR2 signaling reciprocally regulate CD25^lo^AREG^pos^ Treg and CD25^hi^CXCR6^pos^ Treg differentiation by modulating glycolysis

We next examined the regulation of the CD25^lo^AREG^pos^ Tregs, which is uniquely enriched in RA synovial tissue (**Fig. 1, 4a**). *AREG* expression was restricted to Tregs from RA synovial tissues and was absent in blood or fluid (**Extended Data Fig. 4a**). Compared to other Treg subsets, CD25^lo^AREG^pos^ Tregs exhibited lower levels of suppression-associated molecules (**Fig. 3a**). To directly assess the suppressive capacity of CD25^lo^AREG^pos^ Tregs, we sorted total CD25^lo^CXCR6^neg^ Tregs (representing primarily CD25^lo^AREG^pos^ Tregs based on scRNAseq proportions) and CD25^hi^CXCR6^pos^ Tregs from synovial tissue (**Extended Data Fig. 4b**). CD25^lo^CXCR6^neg^ Tregs demonstrated reduced suppressive capacity compared to CD25^hi^CXCR6^pos^ Tregs, consistent with their transcriptional profile **(Fig. 4b)**.

**Figure 4.**
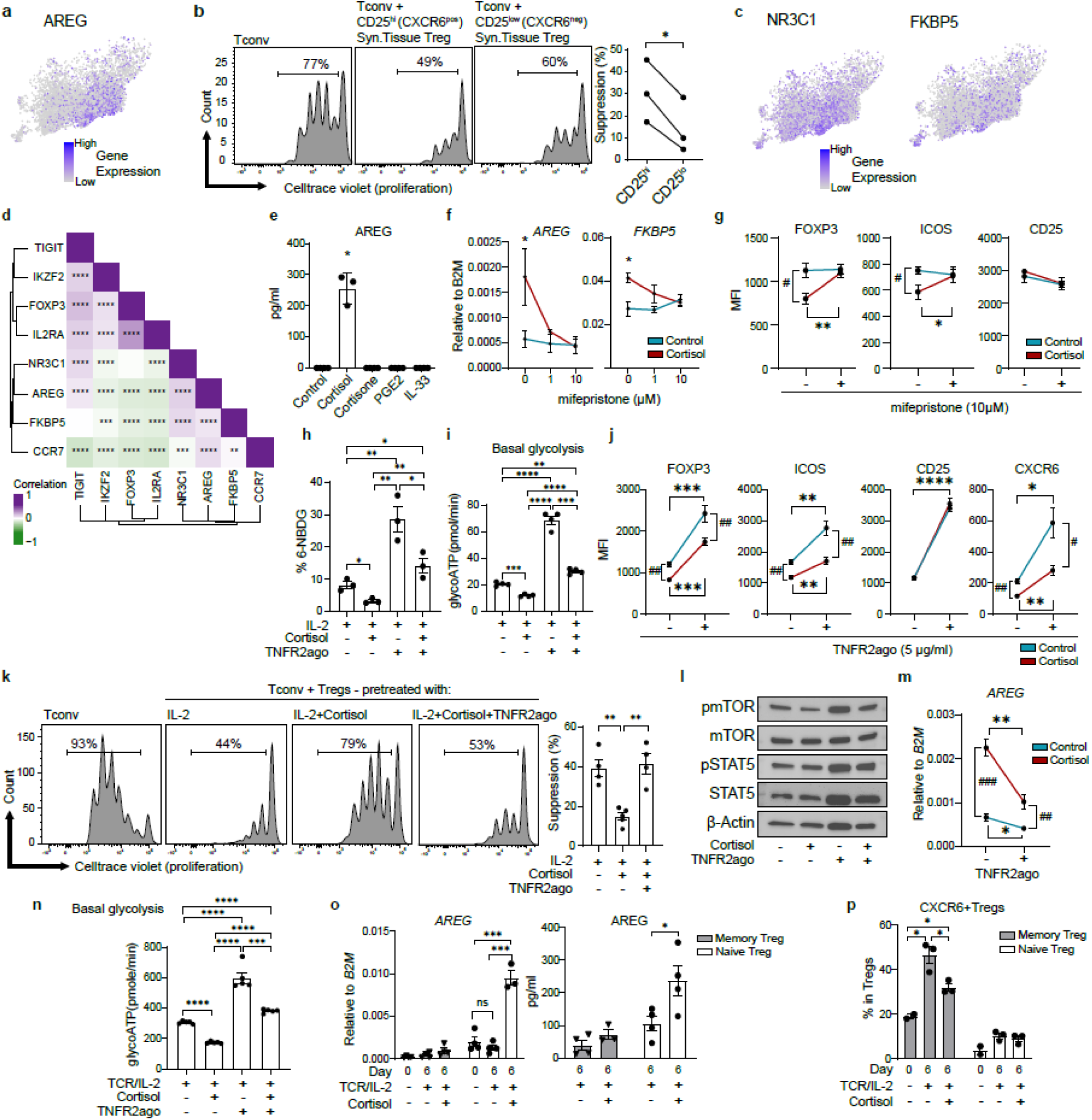
Glucocorticoid and TNFR2 signaling pathway reciprocally regulate CD25^lo^AREG^pos^ Treg and CD25^hi^CXCR6^pos^ Treg differentiation by modulating glycolysis. **a,** UMAP plots highlighting AREG expression in RA synovial tissue. **b,** Suppressive assay of sorted CD25^lo^CXCR6^neg^ Tregs (including CD25^lo^AREG^pos^ Tregs), CD25^hi^CXCR6^pos^ Tregs, and Tconv from RA synovial tissues (n=3 donors). Left: representative histogram; right: mean suppressive capacity. Treg suppressive capacity was calculated as Fig. 3c. **c,** UMAP plots showing *NR3C1*, and *FKBP5* expression in RA synovial tissue Tregs. **d,** Pearson’s correlations between CD25^lo^AREG^pos^ Treg markers and suppressive genes across Treg subsets. Color ranges from yellow (lowest negative correlation) to purple (highest positive correlation), data was trimmed to [−0.6,0.6]. **e-f**, Effects of cortisol on AREG expression in Tregs. ELISA (e) and qPCR (f) analyses shown (n=4 donors). **g,** Flow cytometric analysis of FOXP3 and ICOS in Tregs treated with IL-2 ± cortisol (n=4 donors). **h-l**, Effects of TNFR2 agonism on glycolysis and Treg suppressive function after cortisol treatment. Tregs were isolated from healthy donor blood and pre-treated with IL-2 and cortisol (5μM) or TNFR2ago(5μg/ml) for 3 days (n=4 donors). (h) Flow cytometric analysis of 6-NBDG uptake. (i) Basal glycoATP production (ATP derived from glycolysis as measured by ECAR). (j) Flow cytometric analysis of Treg markers. (k) in vitro suppressive assay. Representative histogram (left) and mean suppressive capabilities (right) are shown. (l) Immunoblot for p-mTOR, total mTOR, p-STAT5, total STAT5. **m-n**, Effects of TNFR2 agonism on *AREG* expression and glycolysis in Tregs stimulated with TCR/IL-2 and cortisol (n=4 donors). (m) qPCR analysis of *AREG* expression. (n) Basal glycoATP production. **o-p**, Effects of cortisol on differentiation of CD25^lo^AREG^pos^ Tregs and CD25^hi^CXCR6^pos^ Tregs from naïve and memory Treg precursors. Naïve and memory Tregs were isolated from healthy donor blood and stimulated with indicated conditions for 6 days (n=4 donors). (o) qPCR analysis (left) and ELISA (right) analyses of AREG expression. (p) Flow cytometric analysis of CXCR6 expression in Tregs. All gene expressions were normalized to β*2M*. Data represent mean ± SEM. Statistical significance was determined using unpaired t-tests, paired t-test (b) or one-way ANOVA in (e): \**p* < 0.05, \*\**p*<0.01, \*\*\**p* <0.001, \*\*\*\**p* <0.0001.

To further characterize CD25^lo^AREG^pos^ Tregs, we examined genes enriched in this Treg subset. Notably, *NR3C1*, encoding glucocorticoid receptor, was significantly upregulated in CD25^lo^AREG^pos^ Tregs compared to the other subsets (*p* = 0.003, log2 FC = 0.4), as was *FKBP5*, a canonical glucocorticoid-responsive gene and regulator of glucocorticoid signaling ^47^ (**Fig. 4c, Supplementary Table 2**). Gene-gene correlation analysis revealed a positive correlation between *AREG*, *NR3C1, FKBP5* and *CCR7*, and a negative correlation with canonical Treg markers such as *IL2RA* and *IKZF2* (**Fig. 4d**). CD25^lo^AREG^pos^ Tregs showed the highest expression of *AREG*, *NR3C1* and *FKBP5* among all non-proliferating Treg subsets (**Extended Data Fig. 4c**), suggesting heightened sensitivity to glucocorticoid signaling in this subset.

To determine whether glucocorticoids could drive CD25^lo^AREG^pos^ Treg differentiation, we treated Tregs with cortisol (active glucocorticoid), cortisone (inactive precursor), IL-33, or PGE2 - the latter two are known to induce AREG in mouse Tregs and human CD4^+^ T cells, respectively ^39,48^ (**Fig. 4e**). Cortisol, but not other treatments, induced AREG expression. This effect was dose-dependently inhibited by mifepristone (**Fig. 4f, left**), a glucocorticoid receptor antagonist, without affecting proliferation (data not shown). Cortisol also increased *FKBP5* expression, further confirming glucocorticoid receptor-dependent transcriptional activation (**Fig. 4f, right**).

To assess whether cortisol impairs Treg function, we analyzed its effect on Treg canonical markers and metabolic features. Cortisol significantly downregulated FOXP3 and ICOS protein levels, and these effects were blocked by mifepristone (**Fig. 4g**), whereas CD25 expression was not affected. Given the known link between glycolysis and FOXP3 stability ^27^ and previous findings that cortisol suppresses glycolysis in effector T cells ^49^, we hypothesized that cortisol-induced Treg dysfunction might involve metabolic suppression. Additionally, because CD25^hi^CXCR6^pos^ Tregs expressed high levels of TNFR2 and exhibited elevated glycolytic activity (**Fig. 3**), - and given that TNFR2 signaling is known to promote glycolysis and Treg stability ^24^-we further hypothesized that TNFR2 signaling might counteract cortisol-induced metabolic suppression. To test these possibilities, we treated Tregs with IL-2 and cortisol, with or without a synthetic TNFR2 agonist (TNFR2ago), and measured glycolytic activity via 6-NBDG uptake (**Fig. 4h**), ATP production from glycolysis (glycoATP) (**Fig. 4i**), and OXPHOS (mitoATP) (**Extended Data Fig. 4f)** using extracellular flux analysis measuring extracellular acidification rate (ECAR) and oxygen consumption rate (OCR) (**Extended Data Fig. 4d-e**). Cortisol significantly suppressed both glycolysis and mitochondrial ATP (mitoATP) production, and both effects were fully restored by TNFR2 stimulation. In parallel, TNFR2 stimulation also restored both FOXP3 and ICOS expression that had been diminished by cortisol (**Fig. 4j**). Notably, TNFR2 stimulation also upregulated CD25 expression regardless of cortisol exposure, and fully reversed the cortisol-mediated downregulation of CXCR6. Consistent with these molecular and glycolytic changes, cortisol significantly impaired the suppressive capacity of Tregs, and TNFR2 stimulation completely restored the suppressive function of cortisol-treated, dysfunctional Tregs (**Fig. 4k**).

Mechanistically, cortisol suppressed mTOR activation without affecting STAT5 activation, consistent with its lack of effect on CD25 expression (**Fig. 4l, Extended Data Fig. 4g**). Further, TNFR2 stimulation selectively restored mTOR activity without affecting STAT5 activation, indicating that TNFR2 supports Treg metabolic function independently of IL-2 signaling. To further investigate the role of TNFR2 in CD25^lo^AREG^pos^ Treg differentiation, we stimulated Tregs with IL-2 and TCR in the presence of TNFR2 agonist. Under these conditions, TNFR2 stimulation markedly downregulated *AREG* expression (**Fig. 4m**). GlycoATP production in Tregs stimulated with IL-2 and TCR (**Fig. 4n**) was significantly higher than in IL-2-treated Tregs (**Fig. 4i**) and was further enhanced by TNFR2 stimulation (**Fig. 4n**), highlighting the synergistic effect of IL-2 and TNFR2 signaling in promoting glycolysis. Cortisol substantially suppressed glycolysis, while TNFR2 stimulation fully reversed this suppression (**Fig. 4n, Extended Data Fig. 4h**). Similarly, mitoATP production, likely reflecting glycolysis-driven OXPHOS, was suppressed by cortisol and restored by TNFR2 stimulation (**Extended Data Fig. 4i-j**). Together, these findings indicate that TNFR2 signaling can effectively counteract the cortisol-induced suppression of glycolysis, restoring both glycolytic activity and suppressive function in Tregs.

Given the opposite effects of cortisol in regulating the phenotype and function of CD25^lo^AREG^pos^ Tregs while reversing the phenotype and function of CD25^hi^CXCR6^pos^ Tregs, we examined whether these subsets arise from different precursor pools. TCR repertoire overlaps, and naïve module scores suggested that CD25^hi^CXCR6^pos^ Tregs may originate from memory Tregs (i.e. CD25^hi^CXCR6^neg^ Tregs and CD25^hi^TNFAIP3^pos^ Tregs), whereas CD25^lo^AREG^pos^ Tregs may derive from naïve Tregs (**Fig. 2**). To test this, we isolated naïve and memory Tregs from healthy donor blood and stimulated them with TCR and IL-2 in the presence or absence of cortisol (**Fig. 4o-p, Extended Data Fig. 4k**). Cortisol induced AREG expression in naïve Tregs at both the gene and protein levels (**Fig. 4o**), while CD25^hi^CXCR6^pos^ Tregs predominantly arose from memory Tregs (**Fig. 4p**). Notably, cortisol significantly downregulated CXCR6 expression in memory Tregs but not in naïve Tregs (**Fig. 4p**). Naïve Tregs expressed higher baseline levels of NR3C1 than memory Tregs, regardless of treatment, whereas FKBP5 was equally induced by cortisol in both subsets (**Extended Data Fig. 4k**).

Together, these results suggest that naïve and memory Tregs respond differently to glucocorticoids and that divergent cytokine and metabolic cues shape the emergence of CD25^lo^AREG^pos^ Tregs versus CD25^hi^CXCR6^pos^ Tregs. Cortisol promotes the differentiation of CD25^lo^AREG^pos^ Tregs from naïve precursors, while TNFR2 signaling supports the differentiation and function of CD25^hi^CXCR6^pos^ Tregs from memory precursors, via upregulating glycolysis and immune regulatory function.

### CD25^hi^CXCR6^pos^ Tregs interact with synovial macrophages which drive their differentiation and function

Given the distinct features of CD25^hi^CXCR6^pos^ Tregs and CD25^lo^AREG^pos^ Tregs, we next explored how these subsets are shaped by the local microenvironment in inflamed RA synovial tissue. We first analyzed which non-Treg cell types co-vary in abundance with Treg subsets in synovial tissue (Dataset A). Correlation analysis revealed that CD25^hi^CXCR6^pos^ Tregs positively correlated with myeloid cells, while CD25^lo^AREG^pos^ Tregs positively correlated with B cells/plasma cells (r=0.3, *p =* 0.006) and showed a positive trend (but not significance) toward correlation with stromal fibroblasts (r= 0.2, *p* = 0.1; **Fig. 5a**).

**Figure 5.**
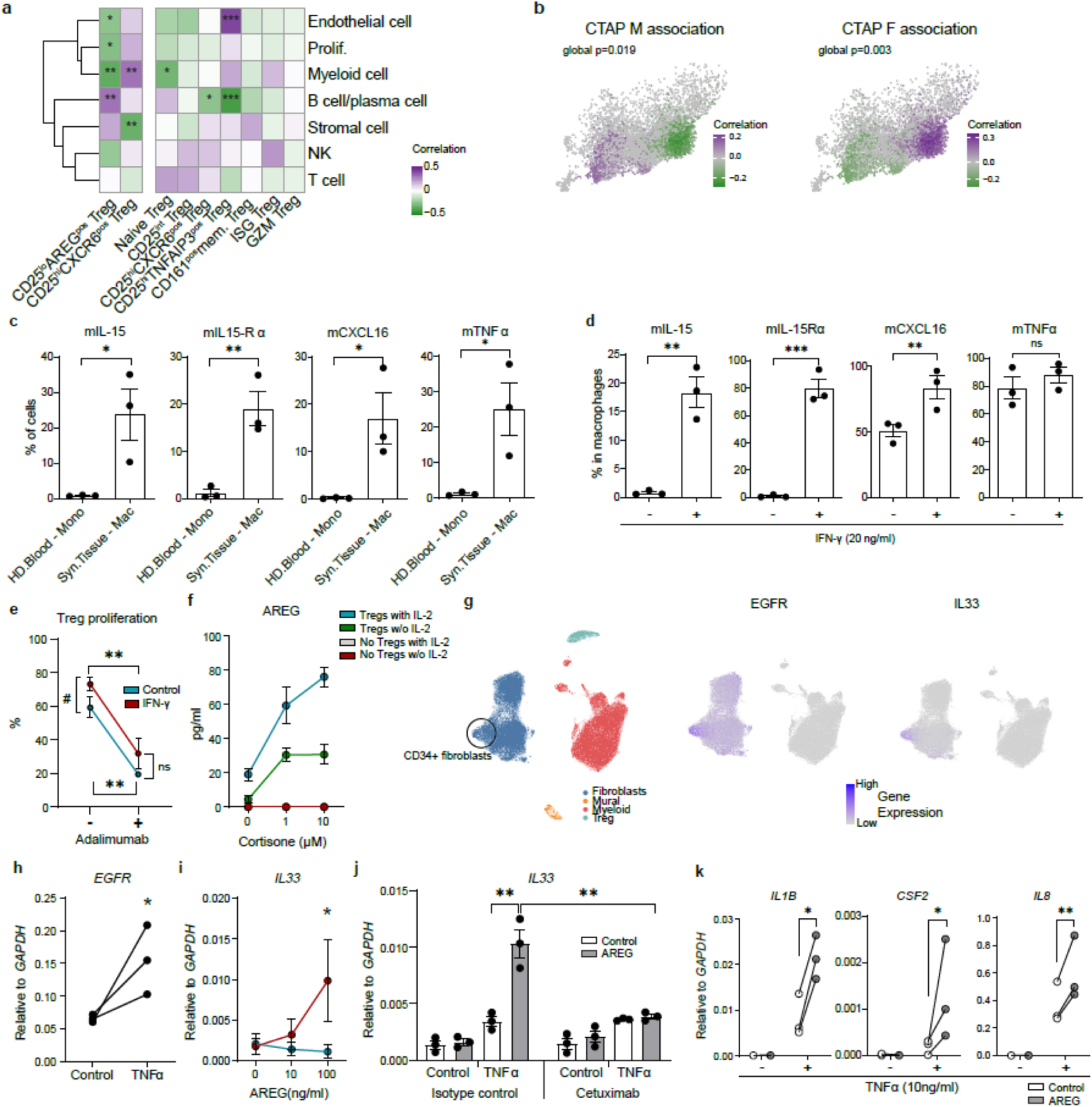
Interaction between Tregs, macrophages, and synovial fibroblasts in RA. **a,** Pearson’s correlation between Treg subset abundance (% of all Tregs, columns) and other cell lineages abundance in the synovial tissue (% of all cells, rows) in dataset A. Color ranges from yellow (lowest negative correlation) to purple (highest positive correlation), data was trimmed to [−0.5,0.5]. **b,** CNA-derived associations between Tregs and fibroblasts enriched CTAPs (left) or myeloid enriched CTAPs (right) (dataset A). Cells are color-coded based on their neighborhood association with the tested CTAP group (red: high correlation; blue: low correlation; Only significantly correlated cells are colored red/blue [FDR < 0.1]). **c-d,** Flow cytometric analysis of membrane bound IL-15 (mIL-15), mIL-15Rα, mCXCL16, and mTNFα in macrophages from RA synovial tissues and monocytes from healthy donor blood (c) and in vitro derived macrophages (d) (n=3-4 donors). **e,** Proliferation of Tregs co-cultured with in vitro-derived macrophage± adalimumab (10μg/ml) or its isotype control for 4 days (n=4 donors). **f,** ELISA measurement of AREG secreted from Tregs-synovial fibroblast co-cultures with cortisone and TCR stimulation ± IL-2 (n=4 donors). **g,** UMAP plot of stromal (fibroblasts, mural), myeloid and Treg lineages from synovial tissue (dataset A). UMAP colored by cell lineage (left), *EGFR* expression level (center), and *IL33* expression level (right). **h-k**, Effects of AREG on synovial fibroblasts. qPCR analysis of *EGFR* expression (h), *IL33* (i-j), and pro-inflammatory genes (k) in synovial fibroblasts (n=3 donors). All gene expressions were normalized to *GAPDH* expression. Data represent mean ± SEM. Statistical significance was determined using unpaired t-tests, one-way ANOVA (f) or ratio paired t-tests (h, i). (\**p* < 0.05, \*\**p*<0.01, \*\*\**p* <0.001, \*\*\*\**p* <0.0001).

To contextualize these findings within the synovial microenvironment, we used the Cell Type Abundance Phenotype (CTAP) classification, which stratifies RA synovial tissue into dominant cell-type-based phenotypes: for example, CTAP-M (myeloid-rich), CTAP-F (fibroblast-rich), and CTAP-TF (T cell and fibroblast-rich) ^32^. Although most Treg clusters were present in donors across different CTAPs, CD25^hi^CXCR6^pos^ Tregs were less abundant in CTAP-F and CTAP-TF donors (**Extended Data Fig. 5a**). Based on donor CTAP classification, we performed co-varying neighborhood analysis (CNA) ^50^ to determine the association between cell neighborhoods in the 2D UMAP space and donor phenotype (e.g. CTAP), in a manner that is independent of cell clusters. We observed a significant enrichment of CD25^hi^CXCR6^pos^ Tregs in myeloid rich CTAPs (M, TM) with depletion of CD25^lo^AREG^pos^ Tregs (*global p* = 0.019, **Fig. 5b, left**). Conversely, CD25^lo^AREG^pos^ Tregs were almost exclusively found in fibroblast rich CTAPs (F, TF), while CD25^hi^CXCR6^pos^ Tregs were less abundant (*global p* = 0.03, **Fig. 5b, right**). Despite the positive correlation between B cells/plasma cells and CD25^lo^AREG^pos^Tregs (**Fig. 5a**), no significant association was observed between Tregs and B cell rich CTAP using CNA (data not shown). Based on these results we tested the possible interactions between these two Treg subsets and myeloid or stromal cells.

First, we focused on the interactions between macrophages and CD25^hi^CXCR6^pos^ Tregs in RA synovium. Monocytes and macrophages can express membrane-bound CXCL16 (mCXCL16), the ligand for CXCR6, as well as IL-15 via IL-15Rα trans-presentation ^51^ and mTNFα ^52^. Building on our prior findings showing that IL-2/IL-15 and TCR signaling promoted CD25^hi^CXCR6^pos^ Treg differentiation (**Fig. 3**), and that TNFR2 stimulation enhanced their suppressive function (**Fig. 4**), we analyzed expression of *IL15*, *IL15RA*, *CXCL16,* and *TNF* in fibroblasts and myeloid cells in synovial tissue (**Extended Data Fig. 5b**). While *IL15* and *IL15RA* were expressed in both synovial fibroblasts and myeloid cells, *CXCL16* and *TNF* were restricted to myeloid cells. At the protein level, synovial macrophages showed higher expression of membrane bound IL-15Rα (mIL-15Rα), mIL-15, mCXCL16, and mTNFα than blood monocytes from healthy donors (**Fig. 5c**). Similarly, blood monocyte-derived macrophages upregulated mIL-15, mIL-15Rα, and mCXCL16 in response to IFN-γ, while mTNFα was constitutively expressed (**Fig. 5d, Extended Data Fig. 5c**). In co-cultures, IFN-γ-stimulated macrophages enhanced Treg proliferation, which was significantly suppressed by blocking mTNFα with Adalimumab, a TNFα neutralizing antibody (**Fig. 5e**). CXCR6 expression remained stable across conditions, and AREG was not detected, likely due to the absence of cortisol in these cultures (data not shown). Together, these data indicate that macrophage-derived mIL-15Rα/IL-15 and mTNFα are essential for CD25^hi^CXCR6^pos^ Treg homeostasis in IL-2-limited RA synovial environment, potentially contributing to immune regulation and disease modulation.

### CD25^lo^AREG^pos^ Tregs interact with synovial fibroblasts which drive their differentiation and function

In parallel, we examined CD25^lo^AREG^pos^ Tregs, which correlated with fibroblasts and B cells (**Fig. 5a**), and were enriched in fibroblast-rich CTAPs (CTAP-F and CTAP-TF) (**Fig. 5b, right**). Given that cortisol induces AREG expression in Tregs (**Fig. 4**), we hypothesized that synovial fibroblasts promote CD25^lo^AREG^pos^ Treg differentiation via glucocorticoid signaling.

RA synovial fibroblasts upregulate hydroxysteroid 11-beta dehydrogenase 1 (encoded by *HSD11B1*) enabling them to convert inactive endogenous cortisone to active cortisol in response to TNFα ^53,54^. We confirmed broad *HSD11B1* expression in fibroblasts in RA synovial tissues (**Extended Data Fig. 5d**) and verified that *in vitro* TNFα-stimulated RA synovial fibroblasts efficiently converted cortisone to cortisol (**Extended Data Fig. 5e**). In co-cultures, Tregs incubated with TNFα-stimulated synovial fibroblasts and cortisone showed increased AREG protein expression, which was further enhanced by IL-2, indicating that both IL-2 and cortisol are important for AREG expression in human Tregs (**Fig. 5f, Extended Data Fig. 5f**). These findings suggest that fibroblast-derived cortisol promotes AREG expression in Tregs, potentially driving the differentiation of dysfunctional CD25^lo^AREG^pos^ Tregs in the RA synovial tissue.

In murine studies, AREG is well characterized for its interaction with epithelial cells and fibroblasts expressing EGFR across multiple organs ^39–42,55,56^. However, the synovium lacks epithelial cells, prompting us to examine transcriptomes of RA synovial fibroblasts (dataset A) (**Fig. 5g**). While *EGFR* was broadly expressed at low levels across all fibroblast clusters, CD34^+^ fibroblasts, previously defined by Zhang et al ^32^, exhibited higher levels of *EGFR* expression (**Fig. 5g, left, center**). This subset also expressed elevated level of *IL33*, a pleiotropic cytokine with both pro- and anti-inflammatory effects depending on the context, tissue, and disease state ^57^ (**Fig. 5g, right**). This observation suggests that AREG may induce IL-33 expression in RA synovial fibroblasts via EGFR. Consistent with our previous report ^58^, we observed TNFα-mediated *EGFR* upregulation (**Fig. 5h**). Here, we found that AREG induced *IL33* expression in a dose dependent manner *in-vitro,* but only in TNFα-stimulated synovial fibroblasts (**Fig. 5i**). Corroborating the transcriptomic data, IL-33 protein levels were significantly increased in AREG-stimulated, TNFα-stimulated synovial fibroblast lysates (**Extended Data Fig. 5g**). However, secreted IL-33 was not detected (data not shown), suggesting that AREG does not trigger cell damage or passive release of IL-33. To confirm the receptor dependency of this effect, we treated synovial fibroblasts with cetuximab, an anti-EGFR monoclonal antibody, and found that this abrogated AREG-induced IL33 expression, demonstrating that the response is EGFR-dependent (**Fig. 5j**). Beyond IL-33, AREG also upregulated inflammatory cytokines in fibroblasts, including *IL1*β, *CSF2* and *IL8* (**Fig. 5k**). However, AREG did not affect *IL6* expression and even inhibited *IL6* expression under IL-17 treated conditions (**Extended Data Fig. 5h, left**). IL-17 did not induce *IL33* expression in synovial fibroblasts, regardless of TNFα stimulation (**Extended Data Fig. 5h, right**). Although AREG has been reported to promote RA synovial fibroblast proliferation ^59^, we did not observe a significant proliferative effect in our system (**Extended Data Fig. 5i**). In contrast, platelet-derived growth factor (PDGF), a well-established fibroblast mitogen^60^, significantly enhanced synovial fibroblast proliferation, regardless of TNFα stimulation.

Together, these findings support a model in which HSD11B1-expressing RA synovial fibroblasts promote the differentiation of dysfunctional CD25^lo^AREG^pos^ Tregs, which in turn secrete AREG and trigger an inflammatory response in synovial fibroblasts, thereby contributing to a pathogenic feedback loop in RA progression.

### Treg phenotypes found in RA synovium exhibit similarities across arthritic diseases

To determine whether the Treg subsets identified in RA synovium are conserved across other autoimmune arthritic diseases, we conducted a comparative analysis of Tregs from juvenile idiopathic arthritis (JIA) (data not yet published, see **Methods**) and psoriatic arthritis (PsA) ^61^ (**Fig. 6a**, **Supplementary Table 1**). We examined Tregs from synovial tissue (JIA only), synovial fluid, and blood (JIA and PsA). After independently analyzing each dataset, we identified 5,523 Tregs from JIA dataset and, 4,096 Tregs from PsA dataset, which were mapped onto the RA Treg UMAP space using the Symphony algorithm, enabling comparison based on shared transcriptomic features ^62^ (**Fig. 6b**).

**Figure 6.**
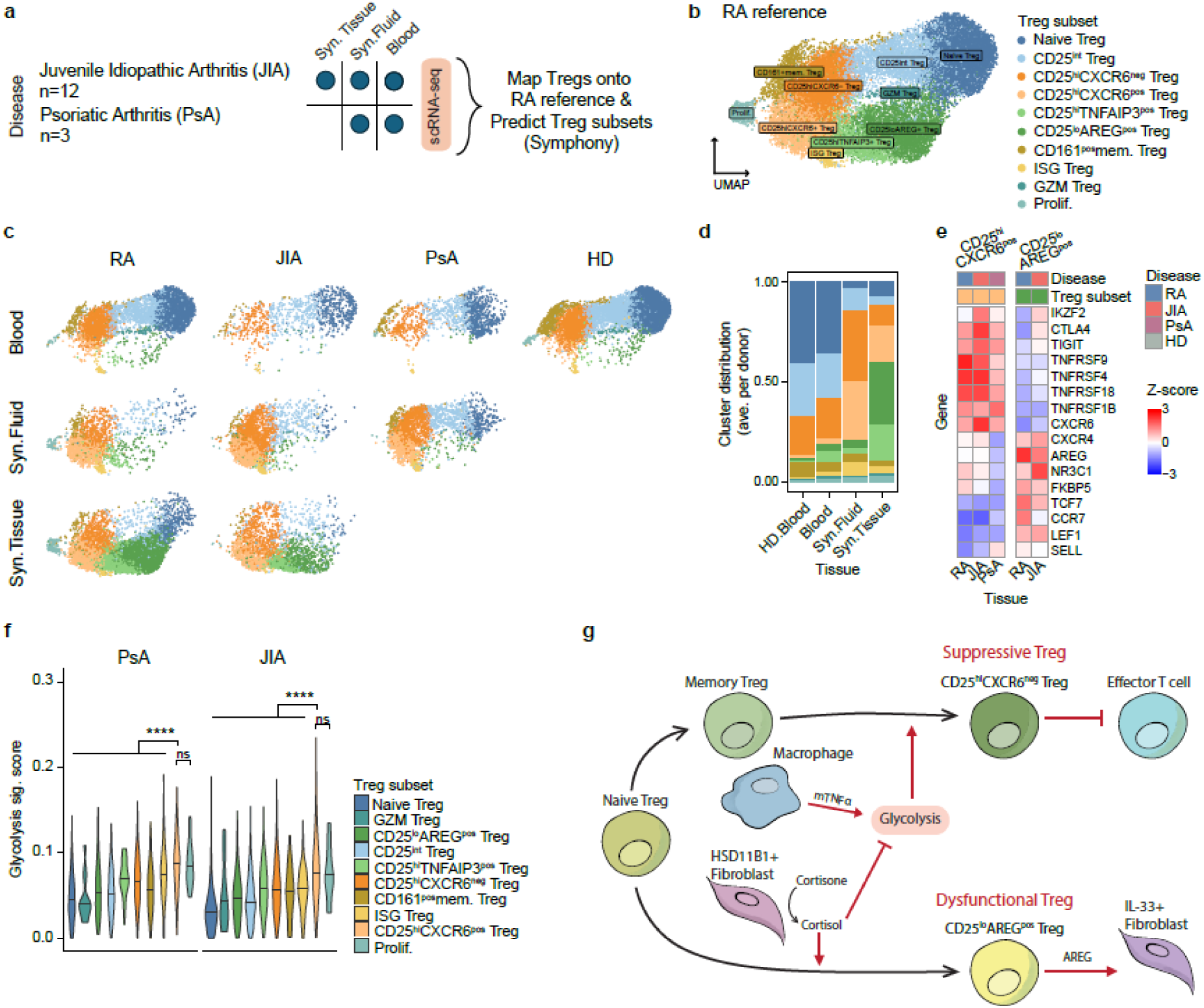
Comparison of RA Treg subsets with Tregs across arthritic diseases. **a,** Schematic overview of the multiple arthritic disease, multi-tissue comparative analysis. scRNAseq datasets from juvenile idiopathic arthritis (JIA) and psoriatic arthritis (PsA) were analyzed separately, and Tregs were independently identified in each dataset. **b,** UMAP plot of RA Treg reference generated using harmony ^79^. Tregs from other arthritic diseases datasets were mapped onto this reference using symphony ^62^, and Treg subset labels were assigned based on transcriptomic similarity. **c,** UMAP plots showing Tregs from RA, JIA, and PsA and healthy donors (HD, columns), split by tissue (rows). **d,** Treg subset proportions across all diseases and tissues, calculated per donor and averaged. **e,** Heatmap of selected genes from pseudobulked samples for CD25^lo^AREG^pos^ Tregs (left) and CD25^hi^CXCR6^pos^ Tregs (right). The CD25^lo^AREG^pos^ Treg heatmap includes only synovial tissue, while CD25^hi^CXCR6^pos^ Treg heatmap includes both synovial tissue and synovial fluid. Z-score values were trimmed to range [−3,3]. **f,** Averaged glycolysis module scores (x-axis) for Treg subsets (colors) across diseases (y-axis). The size of each data point reflects the relative abundance of each subset within total Tregs for that disease. **g,** Schematic summary of the mechanisms described.

Despite differences in disease context and patient demographics, Treg subset distributions in JIA and PsA closely mirrored those observed in RA across all tissue types (**Fig. 6c-d, Extended Data Fig. 6a-b**). In blood, we primarily observed naïve, CD25^int^, CD161^pos^ memory, and CD25^hi^CXCR6^neg^ Tregs, whereas CD25^hi^CXCR6^pos^ Tregs and CD25^lo^AREG^pos^ Tregs were largely absent. In synovial fluid, all three diseases exhibited enrichment of both CD25^hi^CXCR6^neg^ Tregs and CD25^hi^CXCR6^pos^ Tregs. In JIA synovial tissue, we identified both CD25^hi^CXCR6^pos^ Tregs and CD25^lo^AREG^pos^ Tregs, closely matching the RA synovial tissue pattern. PsA dataset lacked synovial tissue, limiting direct comparison to RA and JIA. These findings indicate that shared inflammatory environments in the synovium drive similar Treg heterogeneity across RA, JIA, and PsA.

Further analysis of CD25^lo^AREG^pos^ Tregs in JIA revealed increased expression of *NR3C*1, and *FKBP5*, along with reduced expression of suppressive markers, recapitulating the transcriptional profile observed in RA (**Fig. 6e**). These cells also exhibited elevated expression of naïve Treg-associated genes compared to CD25^hi^CXCR6^pos^ Tregs, supporting the idea that CD25^lo^AREG^pos^ Tregs may derive from naïve precursor and be induced by cortisol in both JIA and RA. Conversely, CD25^hi^CXCR6^pos^ Tregs in JIA and PsA expressed high levels of suppressive markers, similar to their RA counterparts. In RA, this subset is characterized by enhanced proliferative capacity and glycolysis, downstream of IL-2 and TNFR2 signaling (**Fig. 3, 4**). To assess whether this metabolic program is conserved, we calculated glycolysis module scores for Tregs across all diseases (**Fig. 6f**). As expected, CD25^hi^CXCR6^pos^ Tregs consistently showed the highest glycolysis score, followed by other CD25^hi^ Treg subsets. In contrast, CD25^lo^AREG^pos^ Tregs in JIA exhibited low glycolysis, consistent with findings in RA (**Fig. 3, 4**), while naïve Tregs consistently showed the lowest glycolysis score across all diseases. Together, these results support a conserved metabolic and functional profile of CD25^hi^CXCR6^pos^ Tregs and CD25^lo^AREG^pos^ Tregs across multiple inflammatory arthritic diseases. This conservation highlights the role of shared tissue-specific cues in the inflamed synovium in driving both suppressive and dysfunctional Treg states.

In summary, we propose that two distinct Treg subsets emerge in synovial tissue in response to local fibroblasts and macrophages (**Fig. 6g**). CD25^hi^CXCR6^pos^ Tregs maintain high suppressive capacity and are supported by macrophage-derived mTNFα, which enhances glycolysis and Treg stability. In contrast, CD25^lo^AREG^pos^ Tregs arise under the influence of cortisol activated by TNFα-stimulated synovial fibroblasts, and they exhibit impaired suppressive function. These Tregs secrete AREG, which drives pro-inflammatory, IL-33 producing fibroblasts.

## Discussion

Regulatory T cells play a critical role in immunoregulation in both lymphoid and non-lymphoid tissues. Despite their increased frequency in RA synovium, inflammation persists^63^. One explanation for the failure of Tregs to control disease is that local interactions between Tregs and the RA synovial tissue environment might compromise Treg function. Using transcriptomic and functional analyses, we identified two distinct Treg subsets in RA synovial tissues; CD25^hi^CXCR6^pos^ Tregs with high suppressive capacity, and CD25^lo^AREG^pos^ Tregs, which exhibit impaired regulatory function and actively promote inflammatory fibroblast activation.

Our data revealed that macrophages and fibroblasts cooperatively shape the Treg landscape in the RA synovium. Specifically, membrane-bound TNFα (mTNFα) from macrophages supports the homeostasis of suppressive CD25^hi^CXCR6^pos^ Tregs, while fibroblast-derived cortisol promotes the emergence of dysfunctional CD25^lo^AREG^pos^ Tregs. These findings offer mechanistic insight into the paradoxical effects of TNFα-targeted therapies, which have transformed RA treatment but remain ineffective in a significant subset of patients ^64^. We showed that mTNFα blockade on macrophages can impair CD25^hi^CXCR6^pos^ Treg homeostasis, suggesting that TNFα-targeted treatments might inadvertently suppress protective Treg populations – potentially preventing remission or even worsening disease.

Conversely, CD25^lo^AREG^pos^ Tregs, enriched in fibroblast-dominant synovial environments, exhibited diminished suppressive function and release AREG, which promotes IL-33^+^ inflammatory fibroblast activation in TNFα-rich environments. Notably, AREG alone was insufficient to induce this effect, highlighting its role as a context-dependent amplifier of inflammation specifically in TNFα-enriched inflamed tissues. These findings contrast murine studies, where Areg-expressing Tregs are typically tissue-protective and maintain suppressive function. In murine models of lung, muscle, and adipose tissue repair, Areg in Tregs is induced by IL-33 or IL-18 and promotes regeneration without compromising immune regulation ^39,55,56^. However, emerging evidence supports a pathogenic role for Areg in fibrosis across multiple organ ^40–42^. Savage et al. ^41^ demonstrated that Treg-derived Areg promotes liver fibrosis and induces IL-6 production by hepatic fibroblasts, connecting immune regulation to fibrosis and metabolic dysfunction. Our findings extend this concept to human autoimmune tissue, showing that CD25^lo^AREG^pos^ Tregs can reprogram fibroblasts into a pro-inflammatory state in RA. Similarly, macrophage-derived HB-EGF, another EGFR ligand, also was shown to induce IL-33 and promote an invasive fibroblast phenotype in RA ^65^. Together, these data suggest that EGFR signaling in fibroblasts is a central pathogenic axis in human RA, activated by distinct immune-derived EGFR ligands. Other human tissue-adapted Treg populations, such as CCR8□ Tregs in RA and CD161□ Tregs in the gut, have been implicated in tissue repair ^66,67^. However, AREG is not a shared effector among these subsets, emphasizing species-specific and context-specific roles of AREG.

We also demonstrated that cortisol reprograms human Tregs toward a dysfunctional CD25^lo^AREG^pos^ Treg state, which is not recapitulated in murine Tregs. This may reflect differences in glucocorticoid receptor signaling and Treg plasticity across species. In mice, glucocorticoids typically support Treg stability and suppressive function ^68–71^, whereas in our human studies, cortisol downregulated FOXP3, glycolysis, and suppressive capacity. Additionally, cortisol markedly suppressed CXCR6 expression—paralleling findings in murine skin ^72^—and promoted AREG expression specifically in naïve Tregs. These results raise the possibility that glucocorticoid therapy, which is also sometimes part of the therapeutic regimen for RA may disrupt the balance of Treg subsets, favoring dysfunctional CD25^lo^AREG^pos^ Tregs and contributing to persistent inflammation, despite its overall immunosuppressive effects.

Importantly, TNFR2 agonism reversed these cortisol-mediated defects, restoring FOXP3, glycolytic activity, and suppressive function while reducing AREG expression. These findings support the therapeutic potential of TNFR2 agonists—currently in clinical development ^73,74^. However, our findings also raise a cautionary note for Treg enhancement therapies: once therapeutically expanded Tregs enter the inflammatory synovial microenvironment, they may still be susceptible to reprogramming into dysfunctional states, as occurs with endogenous Tregs. As we demonstrated that TNFR2 agonists increases CD25 expression in Tregs in an IL-2/STAT5-independent manner, we suggest that a therapeutic strategy combining TNFR2 agonists and IL-2 may both expand and stabilize functional Tregs, while rescuing or preventing acquisition of the dysfunctional CD25^lo^Treg state in inflamed tissues.

Finally, we show that these Treg states are conserved across inflammatory arthritic diseases, including JIA and PsA. Both CD25^hi^CXCR6^pos^ Tregs and CD25^lo^AREG^pos^ Tregs were identified in these diseases and shared similar transcriptional and metabolic state with their RA counterparts. These findings suggest that inflammatory microenvironments shape a conserved pattern of human Treg states, reinforcing a unifying model of Treg dysregulation across autoimmune arthropathies.

In summary, our study uncovers two divergent Treg states in human RA synovial tissue: one protective, the other dysfunctional and pro-inflammatory. We reveal how macrophage- and fibroblast-derived cues—mTNFα and cortisol—act reciprocally to drive these opposing states, and we demonstrate that AREG functions as a context-dependent amplifier of inflammation. These insights uncover novel mechanisms of Treg destabilization in human autoimmunity and suggest therapeutic strategies. For example, IL-2–based approaches alone may fail due to tissue microenvironment effect, but combining IL-2 with TNFR2agonism may counteract these effects and allow effective functional Treg expansion.

## Methods and materials

### Human research

The study involving Rheumatoid Arthritis (RA) patients was conducted with the approval of the Institutional Review Board at the Brigham and Women’s Hospital. All RA patients included in the study met the American College of Rheumatology (ACR) 2010 classification criteria. Synovial fluid specimens were collected as excess material during routine diagnostic or therapeutic joint aspirations, as prescribed by the patients’ attending rheumatologist. For comparison, blood samples from healthy individuals were acquired from anonymous platelet donors, specifically from the leukoreduction (LR) collars used in blood banking procedures. Patient information **Supplementary Table 3**.

### Isolation of PBMCs from leukoreduction (LR) collars

Peripheral blood mononuclear cells (PBMCs) were isolated LR collars (Specimen Bank, BWH) using Ficoll-Paque (GE Healthcare) density gradient centrifugation. 15 ml of the buffy coat was diluted with 35 ml of ice-cold buffer (PBS with 2 mM EDTA). 25 ml of this diluted suspension was gently added to 15 ml of Ficoll-paque in a 50 ml conical tube. After centrifuging at 400 x g for 40 mins at 20°C, the upper layer was gently removed by aspiration. The mononuclear cell layer was transferred to a new 50 ml conical tube and filled with buffer up to 50 ml. After mixing, cells were centrifuged at 300 x g for 10 mins at 20°C. After removing supernatant, cells were resuspended again with 50 ml of the buffer and centrifuged at 200 x g for 15 mins at 20°C. After repeating this washing step, the number of cells was counted and used for the isolation of human CD4^+^ T cells or cryopreserved in FBS with 10% DMSO for future use.

### Human Tregs culture

Human regulatory T cells were enriched from PBMCs using negative selection with EasySep Human CD4+CD127lowCD25+Regulatory T cell isolation kit (STEMCELL Technologies). The enriched Tregs were subsequently sorted by FACS using a BD FACSAriaIII cell sorter to achieve high purity. The isolated Tregs were stimulated with plate-bound anti-CD3 (5μg/ml, BioLegend) and soluble anti-CD28 (1ug/ml, BioLegend) with human IL-2 (500U/ml), IL-15 (100ng/ml) or plate-bound IL-15/IL-15Rα complex (all cytokines from R&D Systems). Cells were cultured in T cell media consisting of X-VIVO 15 media supplemented with 1mM sodium pyruvate, 1mM L-glutamine, non-essential amino acid, 2-mercapoenthanol, 5mM HEPES and 10% human AB serum for 6 days. For the preparation of the IL-15/IL-15Rα complex, 96 well round bottom plate was coated with recombinant human IL-15Rα (5μg/ml) and incubated overnight at 4°C. The following day, the plate was washed with PBS to remove unbound IL-15Rα, then incubated with recombinant human IL-15 (100ng/ml) for 1 hour at 37°C incubator. Prior to use, the plate was washed with PBS to remove unbound IL-15.

### In vitro human macrophage differentiation

Human CD14^+^ monocytes were isolated from PBMCs by positive selection using CD14 MicroBeads (Miltenyi Biotec) according to manufacturer’s instructions. The isolated cells were cultured in macrophage differentiation medium consisting of RPMI 1640 supplemented with 10% heat-inactivated fetal bovine serum (FBS, Atlanta Biologicals), 1 mM sodium pyruvate, 1 mM L-glutamine, non-essential amino acids, 5mM HEPES (all from LONZA) and 2-mercapoethanol (Sigma-Aldrich). Recombinant human macrophage colony-stimulating factor (M-CSF, 50ng/ml; PeproTech) was added to induce differentiation into macrophages over 3∼4 days. For the Treg-macrophage co-culture experiments, differentiated macrophages were harvested on day 4, and 50,000 cells were seeded into each well of a 96 well round bottom plate. The macrophages were then stimulated with or without human IFN-γ (20ng/ml, R&D Systems) for one day. Following stimulation, the culture medium was replaced with fresh macrophage medium, and the cells were treated with Fc receptor blocking solution (Thermo Fisher Scientific) for 1 hour at 37°C. Subsequently, 100,000 autologous blood Tregs were added to each well containing macrophages, along with soluble anti-CD3 (2ug/ml; BioLegend) to provide TCR stimulation. The co-culture were maintained for 5 days under standard cell culture conditions (37°C, 5% CO2, humidified atmosphere)

### Reverse transcription (RT) and quantitative real-time PCR (qPCR)_qRT-PCR

Total RNA was extracted from cells using the RNeasy Mini Kit (Qiagen) according to the manufacturer’s protocol. The extract RNA was then reverse transcribed into cDNA using the QuantiTech Reverse Transcription kit (Qiagen). For qPCR, gene expression analysis was performed using TaqMan Real-Time PCR assay (Thermo Fisher Scientific). Gene expression levels were normalized to the expression of the housekeeping gene β2-microglobulin (B2M). The relative gene expression was calculated using the comparative threshold cycle (−ΔΔC_T_) method. All experiments were conducted in duplicate. Results are presented as the mean ± standard error of means (SEM) of the biological replicates. Taqman probes used in the study are listed in **Supplementary Table 4**.

### Flow cytometric analysis

For surface staining, cells were incubated with surface antibodies at RT for 30 mins in the dark. Following surface staining, cells were fixed using Foxp3/Transcription factor fixation buffer (eBioscience) at 4°C in the dark for 30 mins or overnight. Fixed cells were then permeabilized using permeabilization buffer (eBioscience) and stained with antibodies targeting intracellular cytokines and transcription factors for 1 hour at RT in the dark. After staining, cells were washed twice with FACS buffer (PBS containing 0.5% BSA and 2mM EDTA) and resuspended in the same buffer for flow cytometric analysis. For phospho-STAT staining, cryopreserved PBMCs were thawed and rested in T cell media for 1 hour at 37°C in a 5% CO2 incubator. PBMCs were then stimulated with various concentrations of IL-2 or IL-15 for 20 mins 37°. Immediately after stimulation, cells were fixed with 4% paraformaldehyde for 15 mins at RT. After washing with PBS, PBMCs were permeabilized with ice-cold 90% methanol for a minimum of 2 hours at −20°C. Following permeabilization, samples were washed with FACS buffer and stained with antibodies targeting phosphorylated STATs for 1 hour at RT in the dark. Data were collected on a BD Fortessa flow cytometer and analyzed using FlowJo software (10.7.1, treestar). The list of fluorescent-conjugated antibodies used for flow cytometric analysis is provided in **Supplementary Table 5**.

### Treg suppressive assay

For the ex-vivo Treg suppressive assay, Tregs and Tconvs were isolated from synovial fluid or tissue by FACS. Tconvs were labeled with CellTrace Violet (Thermo Fisher Scientific) according to the manufacturer’s instructions. 10,000 labeled Tconv cells were co-cultured with an equal number of autologous Tregs at a 1:1 ratio. To assess the effects of cortisol and TNFR2 agonist on Treg function, FACS-sorted Tregs from healthy donor blood were pre-treated with cortisol (5μM) and TNFR2 agonist (5ug/ml) in presence of IL-2 (100ng/ml) for 3 days. On day 4, pre-treated Tregs were washed twice with T cell media to remove residual cortisol and TNFR2 agonist. Freshly isolated Tconv cells from PBMCs were labeled with CellTrace Violet and co-cultured with the pre-treated Tregs. Co-cultures were stimulated with anti-CD3/CD28 Dynabeads (Thermo Fisher Scientific) at various cell-to-bead ratios in T cell media in a 96 well round bottom plate. After 4 days of co-culture, cells were harvested for proliferation analysis by flow cytometry. Treg suppressive capability was calculated using the following formula: Suppression (%) =100 X [(1-(percentage of dividing Tconv with Tregs/ percentage of dividing Tconv without Tregs))

### Measurement of 6-NBDG uptake in Tregs

Tregs were isolated from healthy donor blood using FACS. Sorted Tregs were incubated in glucose free DMEM media (Thermo Fisher Scientific) for 1 hour. Subsequently, 6-NBDG (100μM, Thermo Fisher Scientific) was added and incubated for an additional 45 minutes under standard cell culture conditions. After incubation, cells were washed to remove excessive 6-NBDG, and uptake rate was measured using flow cytometry. For cortisol and TNFR2 agonist-treated Tregs, cells were washed with PBS before proceeding with the 6-NBDG uptake protocol as described above.

### Metabolic assay

For glycolysis (ECAR) and OXPHOS (OCR) measurement, the Seahorse XF T cell metabolism kit (Agilent Technologies) was used. FACS-sorted Tregs from healthy donor blood were treated with cortisol (5μM) and TNFR2 agonist (5ug/ml) in the presence of IL-2 (100ng/ml) or TCR stimulus for 3 days. Tregs (160K cells per well) were seeded in quadruplicate, and the OCR and ECAR were measured using a Seahorse XF96 analyzer (Agilent Technologies) in the presence of the oligomycin A (1.5μM), BAM15(2.5μM) and rotenone/antimycin A (0.5μM each)

### RA synovial fibroblasts culture

Low passage RA synovial fibroblasts were cultured in fibroblast cell culture (FCC) media consisting of DMEM supplemented with 10% fetal bovine serum (FBS, Atlanta Biologicals), 1mM L-glutamine, non-essential amino acids, essential amino acids, 5mM HEPES (all from LONZA), and 0.1mM 2-mercapoethanol (Sigma-Aldrich). Media was exchanged every 3 days, and cells were cultured at 37°C in a humidified incubator with 10% CO2. For the AREG stimulation experiment, 7,000 RA-synovial fibroblasts were seeded in each well of a 96 well flat-bottom plate with FCC media and incubated overnight. On day 2, the media was replaced with low serum FCC media (2% FBS). On day 4, TNF-α (1ng/ml) was added. To test the effects of AREG, varying concentrations (1-100ng/ml) were added on day 5. On day 6, cells were harvested for qPCR analysis. To measure proliferation, 2,000 RA-synovial fibroblasts per well used following the same experimental scheme. Proliferation was quantified using the CyQUANT assay (Thermo Fisher Scientific). To measure the capability for converting cortisone to cortisol, the same experimental design was followed until day 4. On day5, cortisone (1 or 10μM) was added. On day 7, supernatants and cells were collected for cortisol measurement by ELISA (Cayman Chemical), and gene expression using qPCR, respectively.

For the RA-synovial fibroblasts and Treg co-culture experiment, cortisone (1 or 10μM) was added on day 5. On day 7, 100,000 freshly isolated Tregs from blood were added along with ImmunoCult T cell activator (STEMCELL Technologies) and culture for an additional 5 days. On Day 12, the supernatant was collected to measure AREG levels using ELISA (R&D Systems).

### Western blot

The whole Tregs lysate was extracted from Tregs using RIPA buffer (Thermo Fisher Scientific). Protein concentrations were measured using the BCA protein assay (Thermo Fisher Scientific). 10ug of lysates were loaded onto Mini-PRTEAN TGX precast Gels (Bio-rad). After blocking the membrane with 5% Skim milk in Tris-Buffered Saline (TBS), the membrane was probed with primary antibodies specific to target proteins overnight at 4°C. After washing with TBST (TBS with 0.1% Tween-20), the membrane was incubated with secondary antibodies for one hour at RT. After another round of washing with TBST, target protein was detected by ECL solution. Images were taken by CheimiDoc (Bio-rad).

### Synovial tissue processing and cell isolation

Cryopreserved synovial tissue samples were thawed at 37°C in the water bath. The tissue was then washed with RPMI containing 5% FBS using a nylon filter, followed by two additional washed with serum-free RPMI. Subsequently, the synovial tissues were finely chopped using surgical blades and transferred to a FACS tube containing RPMI supplemented with Liberase TL (100μg/ml, Roche) and DNase1 (100μg/ml, Roche). A magnetic stir bar was added to the tube, which was then placed in a beaker containing water at 37°C. The samples were incubated with gentle stirring for 30 minutes. Following incubation, RPMI with 5% FBS was added to the digested tissue. The tissue was then mechanically dissociated using a stainless-steel mesh. The resulting cell suspension was washed with RPMI with 5% FBS and filtered through a nylon mesh to remove any remaining tissue debris. The filtered cell suspension was then used for subsequent experiments.

### Generation of scRNA-seq of synovial fluid and blood from RA patients (Dataset C)

For single-cell RNA sequencing (scRNA-seq), synovial fluid and peripheral blood samples were collected from RA patients (see Supplementary Table 3). Live cells were gated using propidium iodide (PI) and Fixable Viability Dye eFluor™ 780 (Thermo Fisher), followed by isolation of CD45□CD3□CD4□CD25□^/^□CD127^low^ Treg-enriched populations using a BD™ FACSAria III cell sorter (BD Biosciences). A total of 20,000–50,000 sorted cells per sample were stained with cell-hashing antibodies and CITE-seq (TotalSeq-C, BioLegend) antibodies before capture. Cells were then loaded onto a single lane of a Chromium Single Cell Chip (10x Genomics) and processed using the Chromium Single Cell 5′ Library & Gel Bead Kit (10x Genomics), enabling simultaneous capture of gene expression, T cell receptor (TCRα/β) repertoires, and surface protein barcodes. cDNA libraries were generated following the manufacturer’s protocol and sequenced using the Illumina NextSeq 500 platform to a depth of ∼55,000 reads per cell.

### scRNA-seq datasets collection and description

To identify the subpopulations of Tregs in RA tissues, we carried out the main analysis using four datasets (Datasets A-D, Supplementary Table 1). Dataset A (publicly available) included scRNA-seq of whole synovial tissue from 70 RA patients ^32^. Dataset B (publicly available) included scRNA-seq and matched TCR-seq of sorted T cells (CD45^+^CD3^+^) from synovial tissue and PBMCs of 12 donors (shared donors with dataset A cohort) ^75^. Dataset C was newly generated by and included scRNA-seq and matched TCR-seq data of sorted Tregs (CD45^+^CD3^+^CD4^+^CD127^−^) from synovial fluid (n=5) and PBMCs (n=5) of RA patients (total of 9 donors, one donor is shared). Dataset D (publicly available) included scRNA-seq and matched TCR-seq data of sorted Tregs from PBMC (CD4^+^CD25^+^CD127^−^) from 5 healthy donors ^76^.

### Treg isolation from scRNA-seq datasets

We first quantified gene expression counts of datasets B-D from FASTQ files using 10x Genomics CellRanger (10X Genomics, v7.0.1) “count” pipeline ^77^ and the Human reference transcriptome GRCh38 (GENCODE v32/Ensembl 98). For dataset A we obtained the count matrices of the Treg cells as annotated by the original publication ^32^, which were annotated to Human reference GRCh38. Next, we analyzed each dataset independently to identify and extracted Treg cells for downstream analysis. We filtered out low quality cells, keeping those with at least 500 RNA molecules and 250 unique genes and less than 20% mitochondrial reads. In dataset C, donors were marked using hashtag oligonucleotide for which we used “HTODemux” function from Seurat to demultiplex and remove cell from doublet or negative droplets. We performed the following steps as our standard “cluster identification process”. (1) We normalized counts in each cell to 10,000 and log-transformed the data. (2) We selected the top 2000 most highly variable genes (“vst” method) and scaled the normalized expression matrix (to have mean of 0 and standard deviation of 1 for each gene) using Seurat functions ^78^. (3) Principal component analysis (PCA, RunPCA function) followed by Harmony batch correction (v1.2) ^79^ to minimize the variability between donors, with default parameters. For datasets with more than one tissue (B, C) we performed Harmony correction over donor and tissue to also minimize the effect of tissue on downstream analysis. (4) Uniform Manifold Approximation and Projection (UMAP) ^80^, with R uwot package (version 0.1.16) ^81^ with parameters min_dist=0.3, spread=0.8 and ret_extra = ‘fgraph’, to project the cells in two-dimension embeddings. (5) Maximum modularity partitioning of cell-cell similarity graph (“fgraph” from uwot) with Seurat implementation of Louvain clustering algorithm^82^. After initial clustering, for datasets B-D only, we removed doublet cells using scDblFinder algorithm ^83^ with default parameters (this step performed only once per dataset) and repeated steps 1-5 on the remaining singlet cells. (6) To characterize the clusters, we performed differential gene expression analysis using the Wilcoxon rank sum test, implemented in the R Presto package ^84^. We filtered the DEGs (FDR adjusted p < 0.05 and AUC > 0.6).

Following the standard clustering process (steps 1-6) we identified Treg clusters in each dataset as described below. **Dataset A,** included pre-annotated cells from whole synovial tissue from which we used 7440 cells annotated by the original publication as Tregs or proliferating cells. We performed our standard clustering process on these cells and removed contaminating clusters that had marker genes for CD8+ T cells (CD8A, GZMK, NKG7), fibroblasts (COL1A2, COL3A1), B cells (MS4A1, CD74), low quality high mitochondrial gene expressing cells (MALAT1, NEAT1, MT-ND3), and T peripheral/follicular helper cells (Tph/Tfh cells, high CXCL13 gene no FOXP3 expression). **Dataset B,** included **s**orted B and T cells for which we performed our standard clustering process. We first removed B cells clusters based on MS4A1 expression and repeated the standard clustering process. We then removed clusters with CD8 T cells markers (CD8A) or proliferation markers (MKI67) in non CD4+ T cell clusters, and repeated our standard clustering in two iterations. Lastly we called the Treg cluster based on FOXP3 and IL2RA cluster markers. **Dataset C**, included sorted Tregs (CD4+CD127-) this sorting strategy allows for some non Tregs but captures CD25^low^ Tregs which are less commonly studied (most sorted Treg public data is CD4+CD25-). Therefore, after we performed the standard clustering process we removed clusters with less than 20% of their cells expressing FOXP3, we repeated standard clustering process and similar cluster removal two more times to get a clean subset of Tregs. **Dataset D**, included CD4+CD25+CD127-sorted Tregs. We performed the standard clustering process with clustering resolution of 0.8 and removed clusters with less than 20% of the cells expressing FOXP3 or high mitochondrial genes or low quality associated genes in the cluster marker gene list (i.e. MALAT1, MT-ND3), this process performed once.

### Single-cell CITE-seq protein expression

For dataset A, we used the protein expression data from the original publication ^32^. We performed centered-log ratio (CLR) normalization and scaled each protein expression to mean of 0 and standard deviation of 1 across cells within dataset A.

### Treg integrative clustering

We concatenated gene expression count matrices from all Tregs identified in individual datasets (see above) and clustered them using our standard clustering process described in “**Treg isolation from scRNA-seq datasets**” To account for all known potential confounders, we harmonized over donor identity, dataset and tissue, as some datasets included two tissues and some tissues came from more than one dataset. We characterized marker genes overexpression in Treg subpopulations with differential gene expression analysis. To estimate the effect of each Treg cluster on each gene’s pseudo-bulked expression, we fit a generalized linear mixed models (glmm) using lme4 R package (glmer function) ^85^ and extracted differential expression statistics with the Presto R package, GLMM branch (https://github.com/immunogenomics/presto/tree/glmm), following the practice of Korsunsky et al ^86^. We designed the model to account for donor, dataset, and tissue covariates as random effects and the total counts (UMI) for each pseudo-bulked sample. We used the adjusted p-values given by Presto output to determine significance, without additional multiple comparisons adjustment (as described in ^86^), and considered genes with p-value < 0.01 and log2FC > 0.49 (reflects fold-change of 40%) as significant cluster markers.

### Module scores

We show aggregated score of functional groups of genes several times throughout the paper. The rationale for selecting the genes is explained in relevant sections in the results. In all cases, we used “AddModuleScore” function from Seurat to achieve an aggregated score and visualize it in a 2D UMAP space. For metabolic pathways we downloaded gene lists from molecular signatures database (MSigDB) ^44,87^, and filtered them to only include genes that are expressed in at least 100 Tregs (in the integrated Treg dataset). We used “HALLMARK_GLYCOLYSIS” gene set for glycolysis (filtered for 80/200 genes); “HALLMARK_OXIDATIVE_PHOSPHORYLATION” for OXPHOS (filtered for 181 of 200 genes).

### T cell receptor (TCR) sequencing data and analysis

Datasets B-D included paired TCR sequencing data. FASTQ files were demultiplexed (separately from the scRNA-seq FASTQ files) using CellRanger (10X Genomics, v7.0.1). We used VDJ function for assembly and clonotype calling, resulting in a filtered_contig_annotation.csv file that was used for downstream analysis. We used the scRepertoire ^88^ R package to identify TCR sequences obtained for each single cell barcode. We used “filterMulti” argument to filter the data to only include TCR information for: (1) cells in which both α and β chains were captured (2) Top 2 expressed chains per cell in the case of more than two chains captured for one cell. We used cell barcodes to match the TCR information to the cells in the RNA sequencing data. This resulted in clonal information for 88% of cells in dataset B, 90% of cells in dataset C, and 69% cells in dataset D. We used nucleotide sequences from both α and β chains to define the TCR “clone” for each cell. The donor identifier was added to the clone identifier as there is no biological meaning to the unlikely possibility that same clone will be found in two individuals. Clone abundance was later quantified to evaluate clonal expansion, and clones were grouped into five types based on the number of cells they appeared in: Singlet (1 cell), Small (1-3 cells), Medium (4-10 cells), Large (11-30 cells), Hyperexpanded (> 30 cells). Cluster labels were assigned based on matched cell barcodes from the mRNA profiles. We used Morisita-Horn index ^89^ of similarity to calculate shared clones between clusters, as implemented in scRepertoire. The same index was used to plot cluster relationships on the UMAP space.

### Covarying Neighborhood Analysis (CNA)

We used the Covarying Neighborhood Analysis (CNA) ^50^ algorithm to ask if donors with specific Treg sub population have unique synovial tissue composition with higher proportion of specific cell lineages e.g. myeloid, fibroblasts. We used the R implementation rCNA package (version 0.0.99, https://github.com/korsunskylab/rcna) of CNA. To represent global cell type composition of donors in a discrete variable (as required for CNA algoritm), we used a modified version of cell type abundance phenotype (CTAP), first introduced by Zhang et al. ^32^, which divides donors into groups based on the most abundant cell type(s) in their synovium sample. Using dataset A synovial tissue samples only we followed the CTAP classifications from the original paper: F(ibroblasts), T(cells)+F(ibroblasts), M(yeloid), T(cells)+M(myeloid), and T(cells)+B(cells). For simplicity and to increase statistical power given the low number of donors and low number of Tregs per donor, we grouped CTAPs into fibroblasts rich CTAPs (F, T+F), myeloid rich CTAPs (M, T+M), and B cells rich CTAPs (T+B). For each group, we ran a separate CNA analysis to find the cell neighborhoods that are associated with higher abundance of donors with that CTAP. We used the “association.Seurat” function from rCNA with default parameters. The significance of the association was determined by the CNA internal global permutation test with a threshold of p-value < 0.05. For visualization, we used 5% FDR threshold given by CNA and only significantly associated cells were colored based on their correlation level in the UMAP plot shown in figure 5C.

### Single cell RNA data for RA fibroblasts and myeloid cells

We obtained stromal and myeloid cells from dataset A (based on the original paper annotations. Zhang et al. ^32^) and merged them with the subset of Tregs from dataset A as characterized in this paper into one Seurat object. To visualize all these cells in one 2D space, we performed pre-processing as described in steps 1-5 in “scRNA-seq pre-processing” section above. This new UMAP projection is used to show the log counts-per-10K expression of marker genes as described in the results.

### Disease comparison analyses

We downloaded the post CellRanger count matrices of the following publicly available datasets (also detailed in Supplementary Table 1): (1) Juvenile idiopathic arthritis (JIA)-whole tissue scRNAseq data from synovial tissue, synovial fluid and blood of 12 JIA donors. Unpublished data generated by our co-authors, data will be presented by Bolton and Mahony et al. (under review, GSE278962). (2) Psoriatic arthritis (PsA) – scRNAseq of memory enriched CD4 and CD8 T cells from synovial fluid and blood obtained from 3 PsA donors (ArrayExpress accession code <u>E-MTAB-9492</u>) ^61^. We analyzed each dataset independently to isolate the Tregs with the same approach described above for the RA datasets (see “**Treg isolation from scRNA-seq datasets.**”). Next, we mapped each dataset to the harmonized RA Treg reference object with Symphony ^62^ to infer analogous Treg cell states. To generate a reference object from RA Tregs we re-harmonized the RA Tregs in the same way described above and export the results to use by Symphony. We used the Symphony mapQuery function to project external datasets into the RA Treg reference and labeled query Tregs according to their most similar labeled reference Tregs, using the Symphony knnPredict function.

### Statistical analysis

All computational analyses and plotting were carried out using R version 4.2.3 ^90^. Proportional differences in cell populations between tissues were calculated using “scProportionalTest” R package (https://github.com/rpolicastro/scProportionTest), which randomly shuffles cell identities between pairs of samples (in our case, synovium versus blood and synovial tissue versus synovial fluid), finds the log2 proportional differences between the cell numbers in each group and iterating the process to generate p-value and correct for false discovery rate (FDR), as described in Miller et al ^91^. To calculate cell type abundance correlations between Treg sub populations and synovial major cell lineages we used dataset A synovial tissue single cell RNA profiles to calculate the per-donor frequencies of major cell types (as proportion of all cells) and the per-donor frequencies of Treg subpopulations (as proportion of all Tregs). Then we calculated the Pearson correlation between these frequencies across donors.

## Data availability

Raw data for the newly generated dataset C will be deposited in GEO and publicly available upon publication.

## Code availability

All original code will be deposited in GitHub and will be available upon publication.

## Author Contribution

BK, SGO conceived and designed the study. BK, SGO, RS, GD, CM, CB performed experiments and/or data analysis and interpretation. APC, LW provided JIA data and advised on JIA data analysis. LD provided RA synovial tissues. HN advised on fibroblast experiments. SR, IK advised on data analysis. BK, SGO drafted the manuscript. All authors edited and approved the manuscript. DAR and MBB supervised the research.

## Acknowledgement

This work was supported by NIH grants to MBB, P01AI148102. BK was supported by NIH grant (T32AR007530-35). SGO was supported by NIH grant (P01AI148102), the Gordon and Llura Gund Foundation and Israeli Council of Higher Education. RS was supported by Mitsubishi Tanabe Pharma Corporation. LRW is supported by the NIHR Great Ormond Street Biomedical Research Centre. APC was supported by National Institute for Health and Care Research (NIHR) Birmingham Biomedical Research Centre (BRC). CB was supported by Medical Research Council (MRC) Clinical Research Training Fellowship [MR/X001393/1]. LRW was in part supported by the NIHR Biomedical Research Centre at Great Ormond Street Hospital. SR was supported by NIH grants (R01AR063759, P01AI148102, U01HG012009, UC2AR081023). IK was supported by NIH grant (5K01AR078355) and the Gordon and Llura Gund Foundation. DAR was supported by NIH/NIAMS grants (P30 AR070253, R01 AR078769).

---

**Extended Data Fig. 1.**
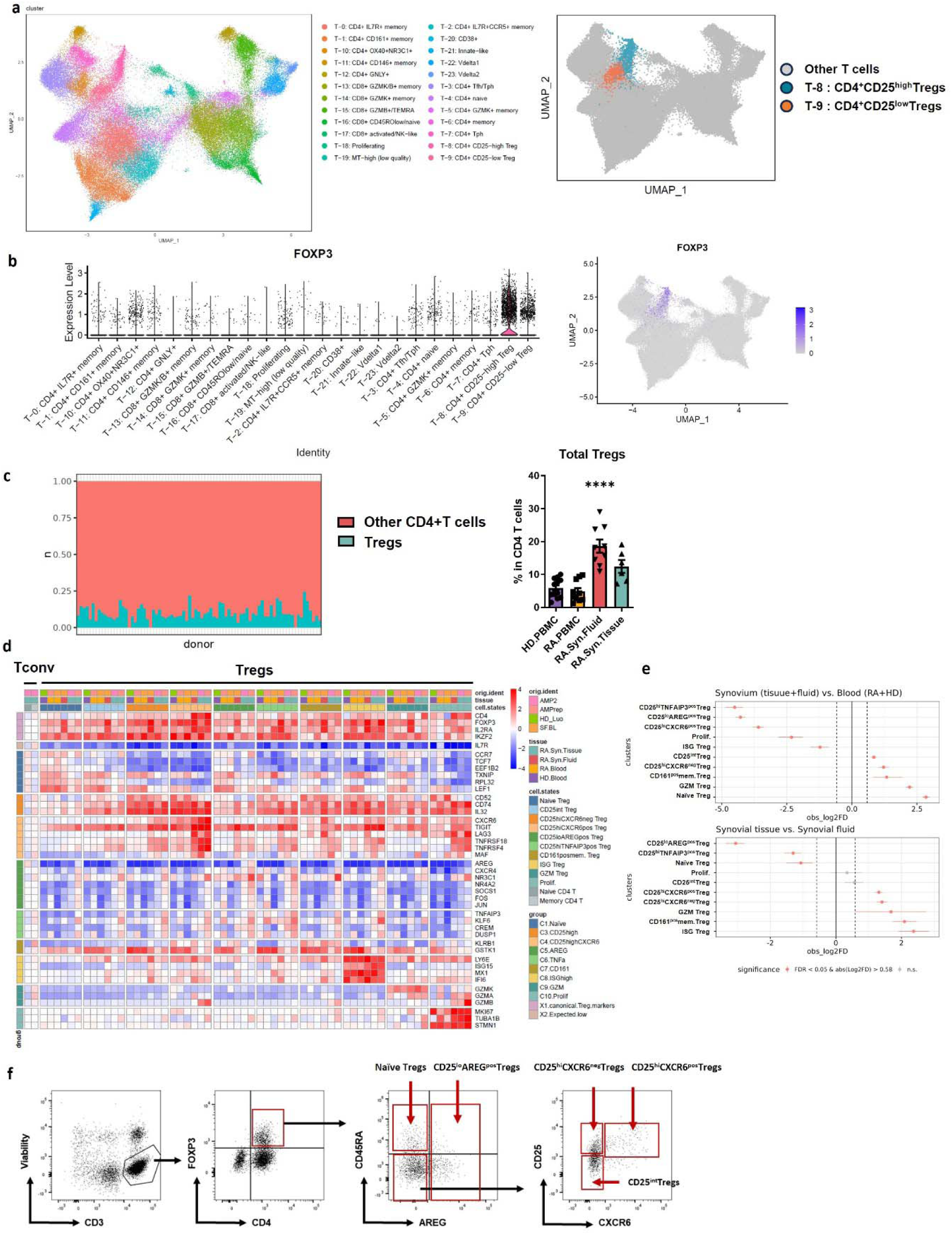
dentification and characterization of Treg subsets across RA tissues using scRNA-seq and flow cytometry. **a,** UMAP plot of T cells from RA synovial tissue, with cluster identities adopted from Zhang et al. ^32^ (left), and corresponding UMAP highlighting Treg subsets within the synovial T cell landscape (right). **b**, Violin plot of log normalized *FOXP3* expression (left) and UMAP plot of *FOXP3* expression (right) across all T cell clusters. **c**, Proportion of Tregs among total T cells per donor in RA synovial tissue (Tregs in teal, other T cells in pink; left). Flow cytometric analysis of the mean percentage of Tregs among total CD4^+^T cells across tissues (right). n=4-10 donors. **d**, Heatmap of pseudobulk expression of selected marker genes in the 10 identified Treg clusters (from all tissues) and in naïve and memory CD4^+^T cells from synovial tissue. Genes were selected based on cluster-specific differential expression and canonical Treg markers. Single-cell counts were summed into pseudobulk samples and log normalized (color bars: top = dataset A–D, middle = tissue, bottom = Treg cluster). Rows are z-scored-normalized relative to non-Treg clusters. Color represents the relative gene expression (blue: low; red: high) **e,** Relative differences in Treg subset proportion between synovium (synovial tissue and fluid) versus blood (RA and HD) and synovial tissue versus synovial fluid. Significant enriched subsets (FDR < 0.05 and mean | log_2_ fold difference | > 1 (permutation test; n=10,000) are shown in red. **f,** Gating strategy of flow cytometric analysis for Treg subsets in RA synovial tissues. Cells were stimulated with PMA and ionomycin in the presence of brefeldin A (BFA) for 2 hours to detect AREG expression. Gating included initial selection by lymphocyte size and granularity, followed by identification of live CD3^+^CD4^+^FOXP3^+^ T cells. Subsequent gates defined Treg subsets.

**Extended Data Fig. 2.**
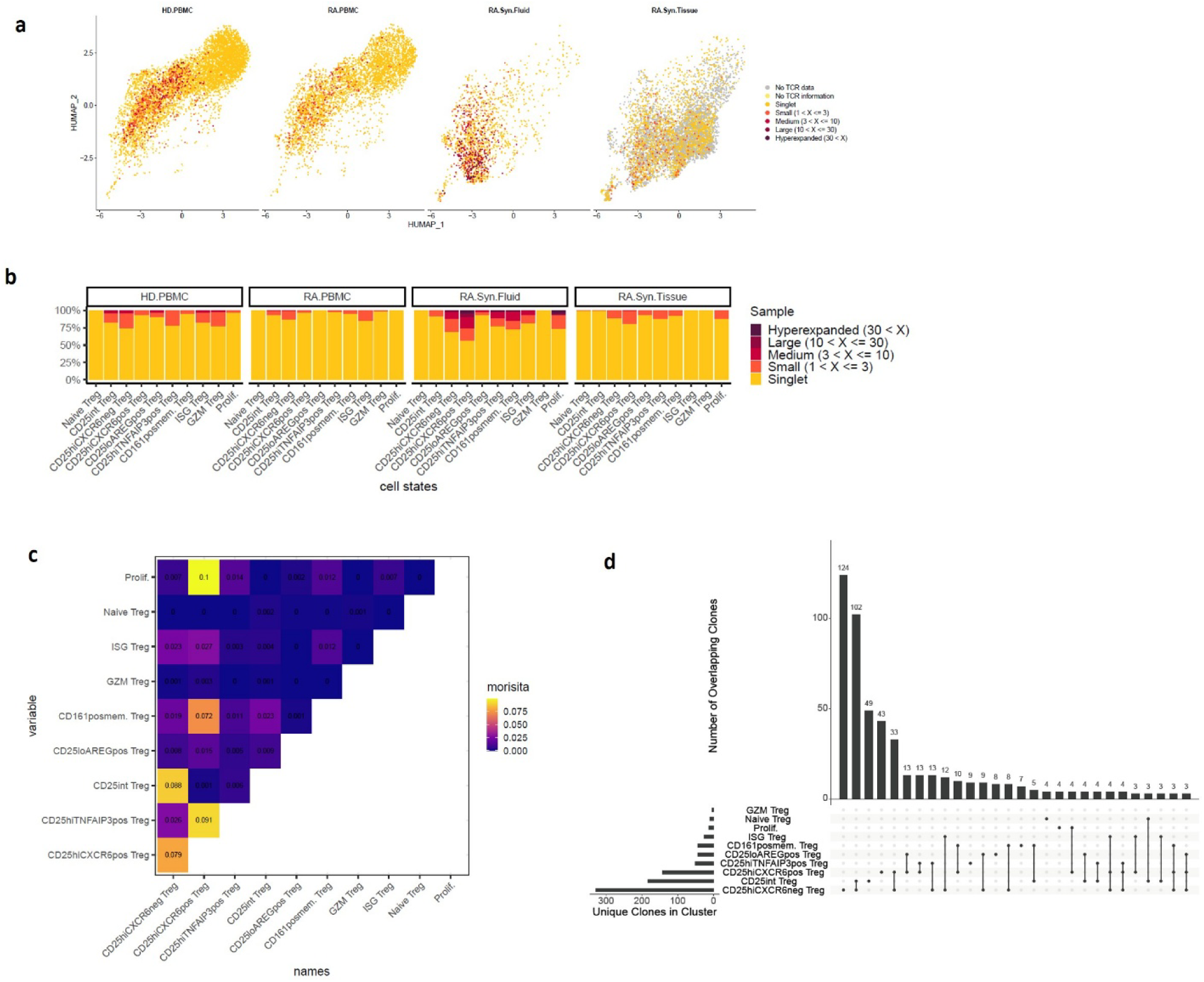
Tissue-specific clonal expansion and inter-cluster overlap of Treg subsets in RA. **a,** Proportions of clonally expanded cells within each Treg cluster, stratified by tissue. Only datasets B-D are shown. Clonal categories are as in Fig. 2a. **b,** Clonal expansion visualized on UMAP plot, split by tissue of origin. Cells are color-coded based on their clonal expansion level, determined by the CDR3 sequences of both α and β chains. “No TCR data” represents cells from dataset A, which lacked scTCRseq data. “No TCR information” represents cells with scRNAseq but were not captured by scTCRseq. Singlets denote cells with unique clones. Expanded clones are categorized as follows: Small (2-3 cells per clone), Medium (4-10 cells per clone), Large (11-30 cells per clone), and Hyperexpanded (>30 cells per clone). **c,** Heatmap of pairwise clonal overlap between all Treg clusters, quantified using Morisita’s index. Higher values indicate greater clonal similarity. **d,** Clonal overlap summary using an UpSet-style plot. The horizontal bar graph (left) shows the number of unique clones per cluster. The vertical bar graph (top) shows the number of shared clones between clusters connected below by dots and lines.

**Extended Data Fig. 3.**
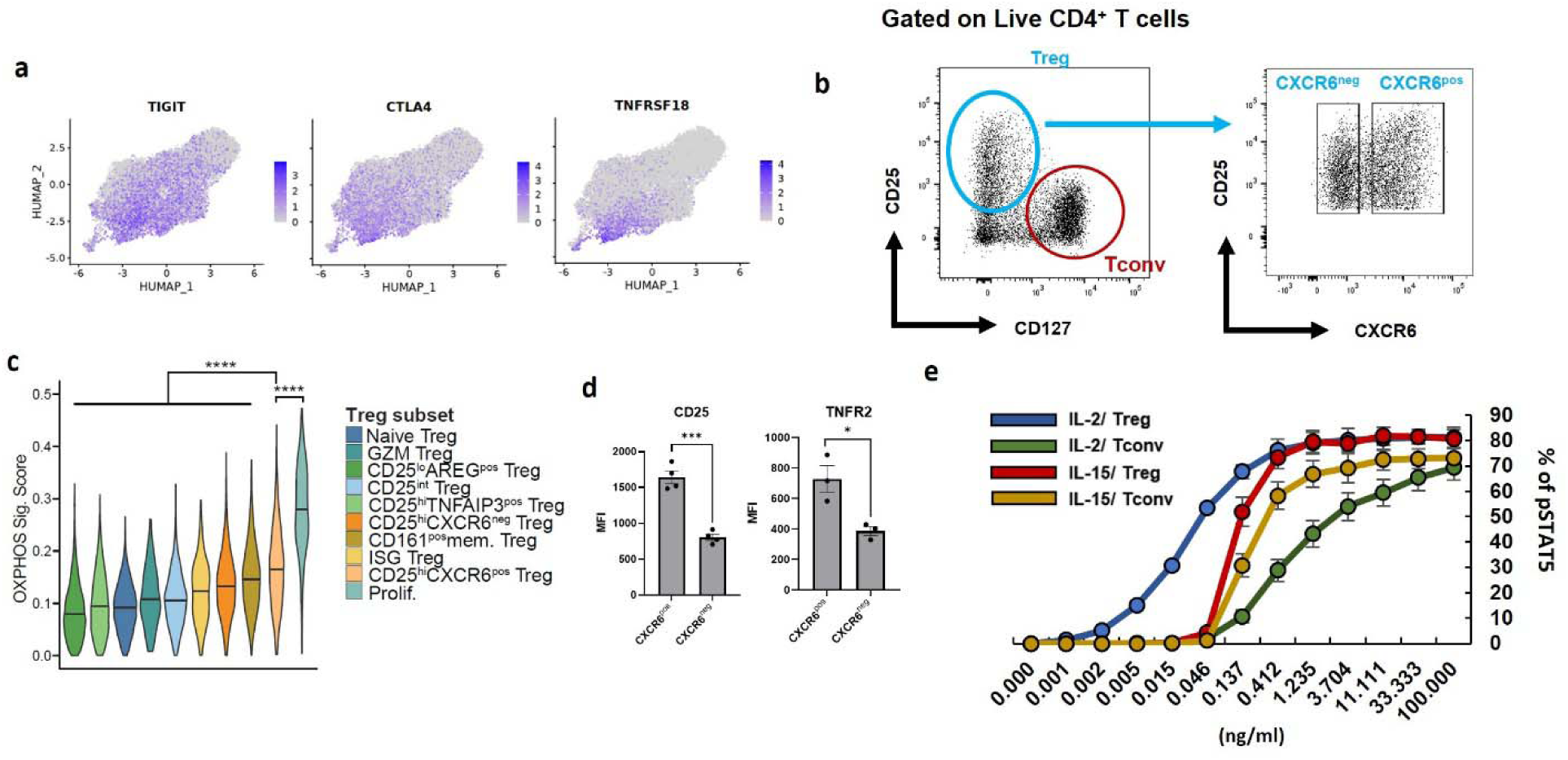
Metabolic and suppressive features of CD25^hi^CXCR6^pos^ Tregs. **a,** UMAP plot colored by expression of *TIGIT*, *CTLA4*, *TNFRSF18*. Grey = no expression; dark purple = highest expression. **b**, Gating strategy for isolating Tregs and Tconv from RA synovial fluid. **c**, OXPHOS module scores (based on 181 genes from the “HALLMARK_OXIDATIVE_PHOSPHORYLATION” gene set, MSigDB ^44,87^) shown as across Treg subsets. The Wilcoxon test was used to assess differences in OXPHOS module score between CD25^hi^CXCR6^pos^ Tregs and each of the other Treg subsets. **d**, Flow cytometric analysis of CD25 and TNFR2 expression in Tregs from synovial tissues (n=4 donors). **e**, Phosphorylated STAT5 in Tregs and Tconv from HD blood after IL-2 or IL-15 treatment (n=4 donors). Data represent mean ± SEM. Statistical significance was determined using unpaired t-tests: \**p* < 0.05, \*\**p*<0.01, \*\*\**p* <0.001, \*\*\*\**p* <0.0001.

**Extended Data Fig. 4.**
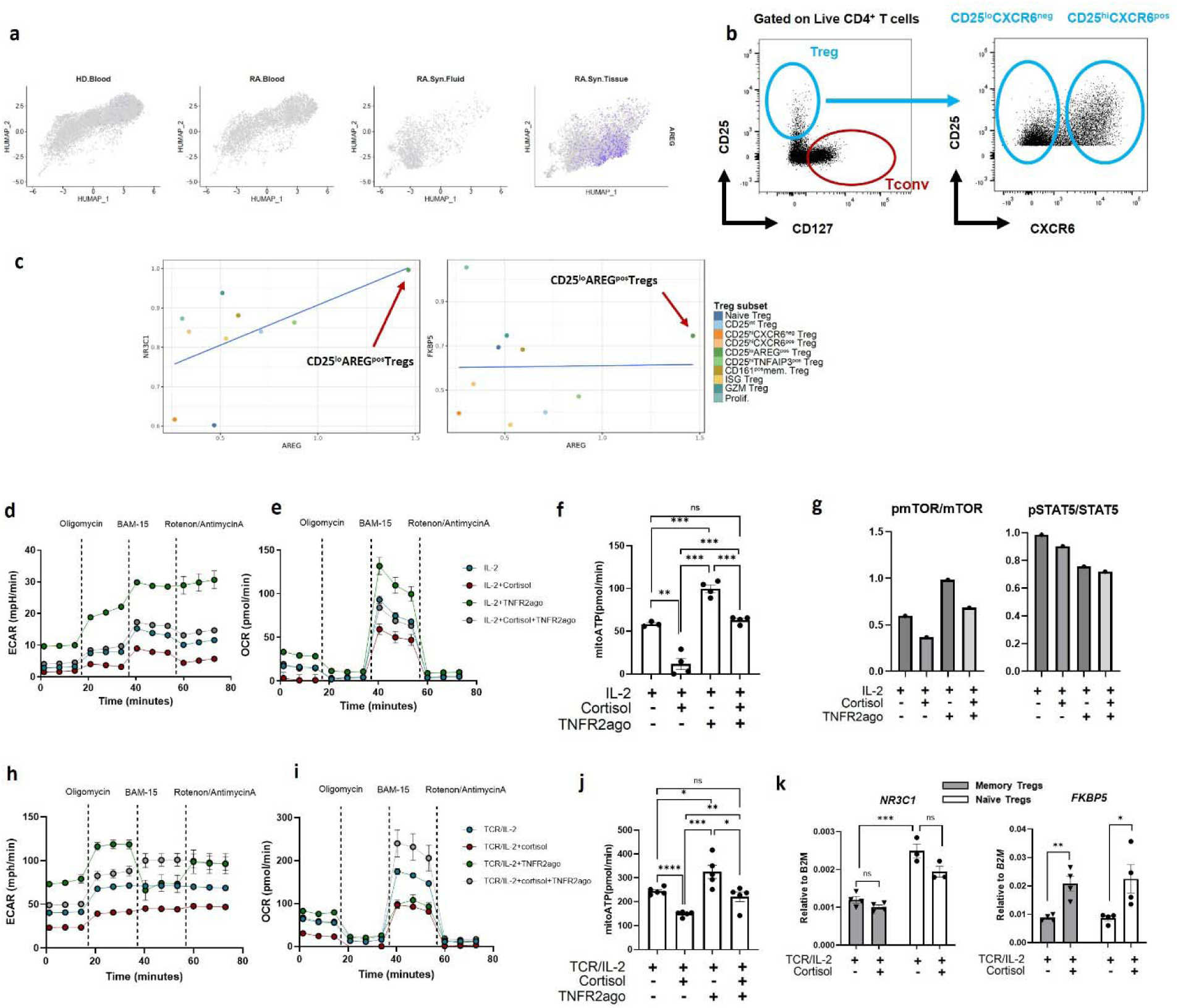
Functional and transcriptional responses to cortisol and TNFR2 signaling. **a**, UMAP plots displaying AREG expression across tissues. **b**, Gating strategy for in vitro Treg suppressive assays. **c**, Pairwise correlation analysis of gene expression in pseudobulked Treg subsets. Each plot shows log-normalized expression comparisons: AREG vs. NR3C1 (left) and AREG vs. FKBP5 (right). Blue regression lines indicate linear fits between the variables. **d-f**, Effects of cortisol and TNFR2 stimulation on glycolysis in Tregs treated with IL-2 and cortisol (5μM) for 3 days (n=4 donors). (d) Extracellular acidification rate (ECAR); (e) oxygen consumption rate (OCR) before and after sequential injection of oligomycin (1.5 μM), BAM15 (2.5 μM), and rotenone/antimycin A (0.5 μM each); (f) basal mitochondrial ATP production rate. **g**. Densitometric quantification of the ratio of phosphorylated to total mTOR (left) and total STAT5 (right), normalized to loading control (β-actin). **h-j**, Effects of cortisol and TNFR2 stimulation on glycolysis in Tregs stimulated with TCR/IL-2 and cortisol (5μM) or TNFR2ago(5μg/ml) for 4 days (n=4 donors). ECAR (h), OCR (i) and basal mitoATP production rate (j) were measured as in (d-f). **k**, qPCR analysis *NR3C1* (left) and *FKBP5* (right) expression of naïve and memory Tregs from healthy blood after 6-day stimulated with TCR/IL-2 under the indicated conditions (n=4 donors). All gene expressions were normalized to β*2M*. Data represent mean ± SEM. Statistical significance was determined using unpaired t-tests: *p < 0.05, **p<0.01, ***p <0.001, ****p <0.0001.

**Extended Data Fig. 5.**
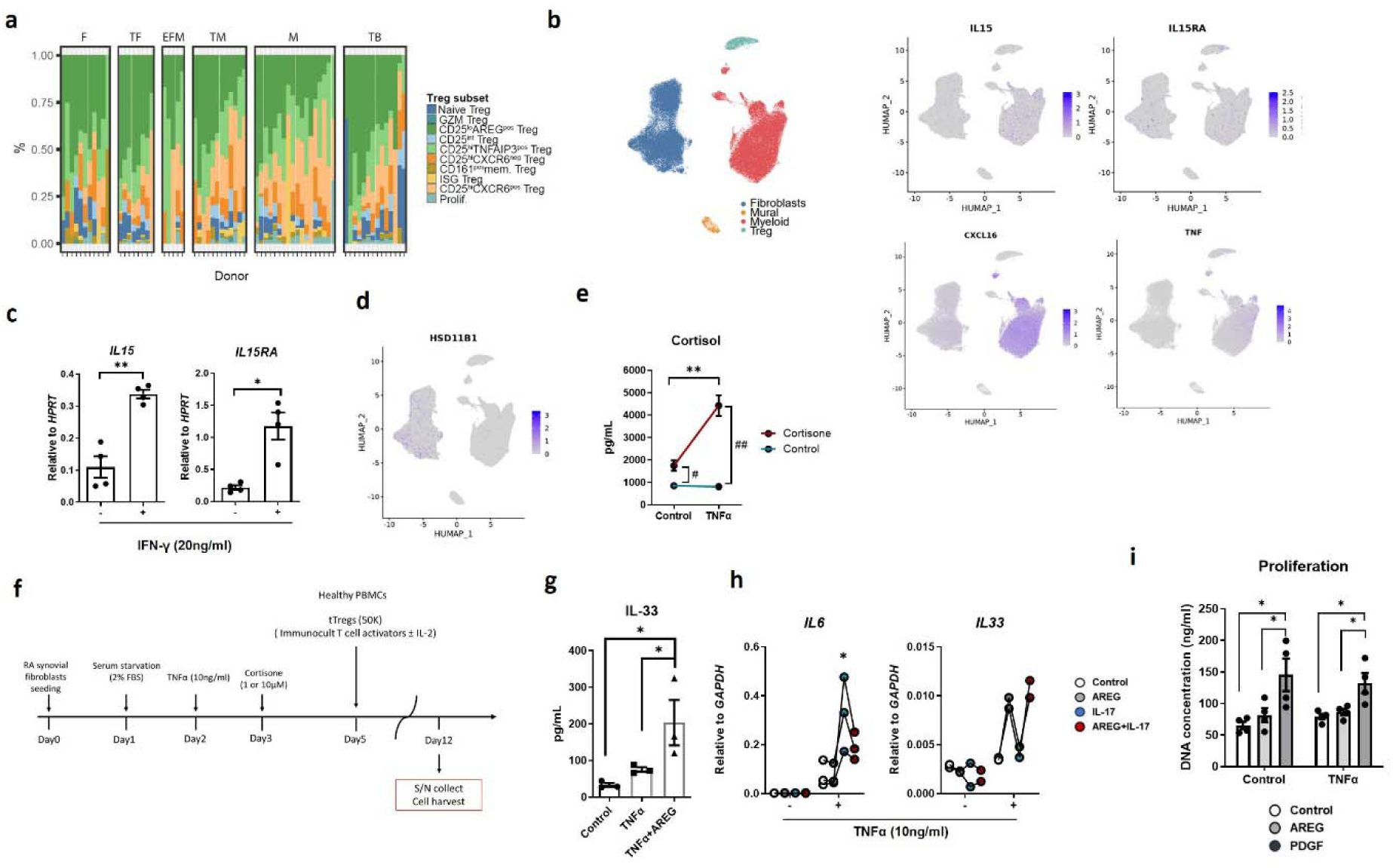
Cell-type–specific pathways influencing Treg states in RA synovium. **a,** Proportions of Treg clusters in individual donors (columns) grouped by CTAP in dataset A. **b,** UMAP plot of stromal (fibroblasts, mural), myeloid and Treg lineages from synovial tissue (Left, dataset A). UMAP colored by cell lineage (left), expression of *IL15*, *IL15RA* (right top), *CXCL16* and *TNF* (right bottom) on UMAP. **c,** qPCR analysis of *IL15RA* and *IL15* expression in vitro generated macrophages (n=4 donors). Gene expressions were normalized to *HPRT* expression. **d,** Expression of *HSD11B1* shown on UMAP. **e**, ELISA of cortisol conversion by RA synovial fibroblasts treated with TNFα (10ng/ml) and cortisone (10μM) for 3 days (n=3 donors). **f,** Schematic of co-culture of Tregs and RA-synovial fibroblasts. **g**, ELISA measurement of IL-33 from RA synovial fibroblast cell lysates (n=3 donors). **h**, qPCR analysis of *IL6* and *Il33* expression in RA synovial fibroblasts stimulated under indicated conditions (n=3 donors). Gene expressions were normalized to *GAPDH* expression. **i,** Proliferation of RA synovial fibroblasts treated with AREG or PDGF, measured by quantifying DNA content (n=4 donors). Data represent mean ± SEM. Statistical significance was determined using unpaired t-tests: \**p* < 0.05, \*\**p*<0.01, \*\*\**p* <0.001, \*\*\*\**p* <0.0001.

**Extended Data Fig. 6.**
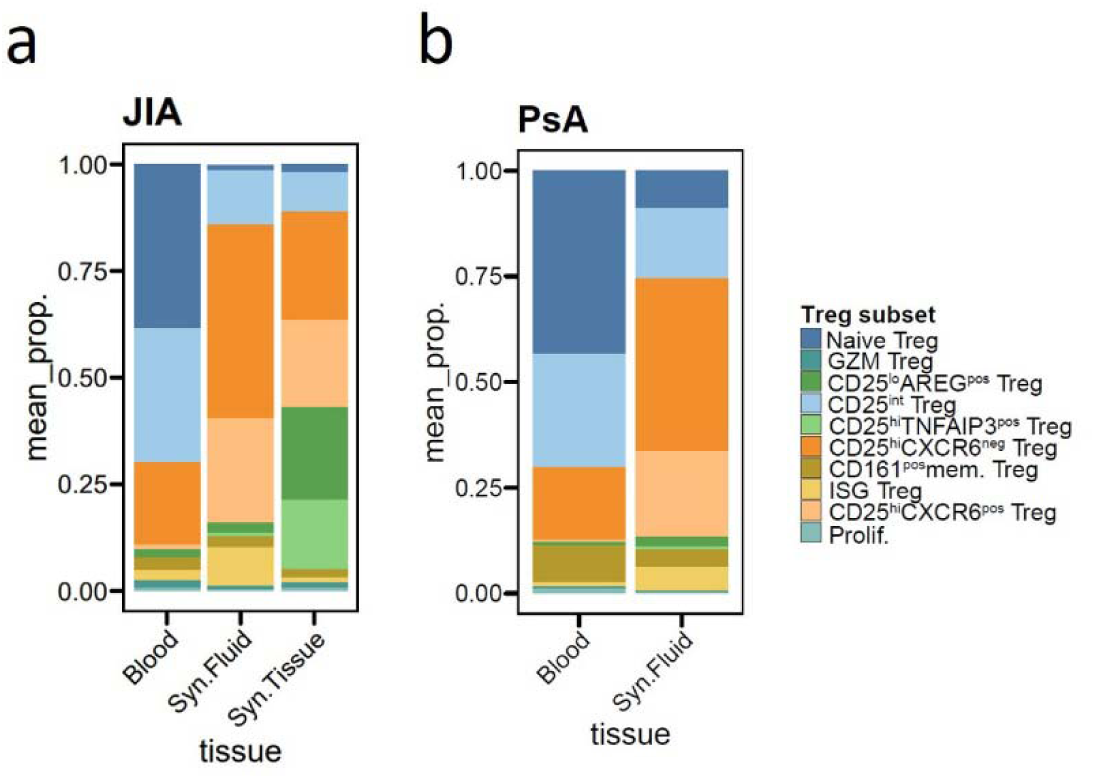
Treg subset distributions across tissues in JIA and PsA. Treg cluster proportions were calculated per donor and averaged for the following: **a,** tissue (blood, synovial fluid and synovial tissue) in JIA; **b,** tissue (blood and synovial fluid) in PsA.

